# Cortical astrocytes flexibly encode reward contingencies and shape conditioned behavior

**DOI:** 10.64898/2025.12.08.693010

**Authors:** Jacqueline E. Paniccia, Roger I. Grant, Annaka M. Westphal, Rachel E. Clarke, Elizabeth M. Doncheck, Bayleigh Pagoota, Jayda Carroll-Deaton, Bogdan Bordieanu, Sophie Buchmaier, Lydia G. Klumb, Lisa M. Green, Joshua A. Boquiren, James M. Otis, Michael D. Scofield

## Abstract

Learned associations between environmental cues and reward drive motivated behavior, yet how specific cell types support this process remains unclear. Using longitudinal two-photon calcium imaging, we tracked dorsal medial prefrontal cortical astrocytes throughout the acquisition, expression, and reversal of Pavlovian sucrose conditioning. As learning progressed, astrocytes exhibited time-locked, spatially coordinated calcium signals that differentiated correct behavioral action from mistakes, evolving from broad outcome encoding to selective representation of responses associated with the reward-conditioned stimulus. Omission testing revealed that prefrontal astrocytes preferentially respond to the cue-reward association, rather than the conditioned stimulus or reward alone. When reward contingencies were reversed, astrocytic activity rapidly adapted to track the new cue-reward association and encode updated and outdated motivated behavioral actions. Finally, astrocytic ablation attenuated motivated behavior during initial associative learning and prevented persistence of conditioned reward seeking when reward contingencies were updated or unpredictable. These findings reveal prefrontal astrocytes are functionally plastic elements that regulate reward-seeking behavior across associative learning.

**Teaser:** Prefrontal astrocytes flexibly encode the cue-reward associations that drive conditioned reward-seeking behavior.

## INTRODUCTION

The ability to adapt behavioral responses based on relevant stimuli is necessary for survival in ever-changing environments. Learned associations between external cues that signal reward availability or those that signal threat shape appropriate behaviors and drive goal-directed, motivated action. The dorsal medial prefrontal cortex (dmPFC) serves as a critical neuroanatomical hub to orchestrate decision-making, learning, executive function, and goal-directed behavior through its excitatory projections to the thalamus, striatum, and ventral tegmental area^1–11^. Accordingly, discrete neuronal ensembles within dmPFC output circuits encode specialized information pertaining to reward-predictive cues, reward delivery, and reward-motivated behavioral action^5,12^. Specifically, several unique activity-based neuronal clusters emerge throughout conditioning in a Pavlovian sucrose task^12^. However, precisely how dmPFC network activity is refined, maintained, and updated across associative learning is less understood.

Recently, astrocytes have emerged as key regulatory elements within neural networks^13–18^. Specifically, cortical astrocytes coordinate neuronal responses, shaping the excitatory-inhibitory balance within the PFC required to direct goal-directed action and response adaptation in reversal learning tasks^14,19–22^. Likely through this ability to influence neuronal activity, astrocytes also facilitate memory stabilization during associative learning tasks^23^. While these findings implicate a role for astrocytes in directing PFC-dependent behavior, precisely how dmPFC astrocytes are engaged throughout reward learning, and if they actively participate in guiding motivated behavior, remains unknown. Astrocytic activity is commonly evaluated by measuring oscillations in intracellular calcium (Ca^2+^)^24–26^, which can be readily quantified through genetically encoded Ca^2+^ indicators *in vivo*^27,28^. These astrocytic Ca^2+^ elevations have been shown to track cue-reward associations and encode reward-related information within the hippocampus and striatum^29,30^, and chemo- or optogenetic stimulation of astrocytes in these regions is sufficient to drive conditioned appetitive^31^ and goal-directed behaviors^30^. Considering that cortical astrocytes can bidirectionally modulate neuronal dynamics to influence cognitive flexibility^22,32,33^, we sought to understand how dmPFC astrocytic activity patterns correspond to appropriately adapting behavior in PFC-dependent reward learning tasks.

Using a head-fixed Pavlovian sucrose conditioning task combined with two-photon microscopy and subsequent astrocytic ablation, we tested the hypothesis that dmPFC astrocytes functionally encode the cue-sucrose association responsible for conditioned reward seeking (anticipatory licks). We visualized cortical astrocytes with single-cell resolution and measured Ca^2+^ transients throughout acquisition of cue-sucrose conditioning, reversal of reward contingencies, and omission of the sucrose-paired cue or sucrose itself. We then lesioned dmPFC astrocytes using the gliotoxin l-α-aminoadipate (L-AA) to uncover whether these cells shape the motivated behavioral actions that emerge with cue-reward associative learning. Collectively, these experiments reveal that prefrontal astrocytes dynamically encode cue-reward associations to regulate conditioned reward seeking across various phases of learning.

## RESULTS

### dmPFC astrocytes track associative learning and encode correct behavioral responses, mistakes, and cued reward delivery

Cortical astrocytes differentiate between types of neurotransmission with distinct cellular Ca^2+^ signatures capable of regulating activity within both the astrocytic syncytia^34,35^ and neuronal network^19,20^ to drive complex behaviors^32^. Here, we combined head-fixed Pavlovian sucrose conditioning with deep-brain two-photon Ca^2+^ imaging to longitudinally measure dmPFC astrocytic activity dynamics during cue-reward associative learning (**Fig. 1A, B**). Head-fixed mice were trained to associate one tone conditioned stimulus (CS+), but not another (CS−), with sucrose delivery. After conditioning, mice increased licking behavior (**Fig. S1A**), displayed anticipatory licking between CS+ onset and sucrose delivery (**Fig. 1C, D**), and appropriately discriminated between cue types (**Fig. 1E; S1B, C**). We then classified the behavioral responses based on the change in lick rate to the CS+ or CS− and found correct licking (high anticipatory licking to the CS+) increased, while incorrect withholding (low anticipatory licking to the CS+) decreased, across conditioning (**Fig. 1F**). Altogether, these results demonstrate mice acquired the cue-sucrose association, updated behavioral responding across conditioning, and displayed anticipatory licking following CS+ presentation.

**Figure 1.**
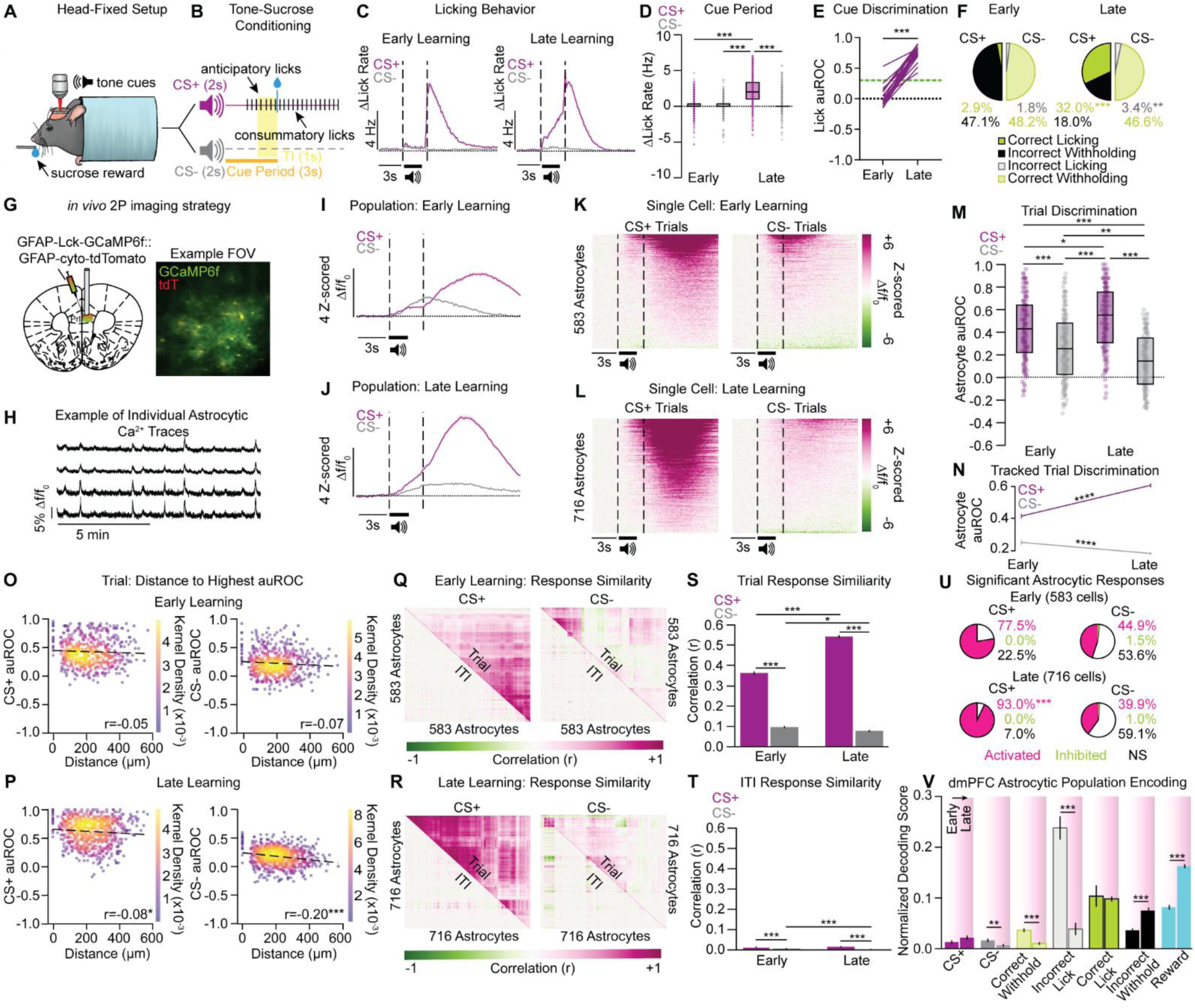
dmPFC astrocytic Ca^2+^ activity tracks cue-reward associative learning and encodes reward-conditioned stimuli, cued reward delivery, and correct and incorrect behavioral responses. **A, B)** Head-fixed set-up **(A)** for *in vivo* Ca^2+^ imaging during Pavlovian sucrose conditioning **(B). C)** Averaged Δ lick rate traces show an increase in anticipatory licks following CS+ exposure from early to late conditioning (n=20 mice). **D)** The change in anticipatory lick rate significantly increased during the cue period of CS+ trials from early to late learning, with no change in lick rate during the cue period of CS− trials (n=20 mice, 21-22 FOVs; Day x Trial Type interaction: F_1,4296_=921.45, *p*<0.001). **E)** Behavioral discrimination (anticipatory licking to the CS+ vs CS−) significantly increased from early learning to late learning (T_19_=17.95, *p*<0.001). Green line indicates auROC threshold of 0.3. **F)** Correct licking (high anticipatory lick rate during the CS+ cue period) significantly increased from early to late learning (Χ^2^(1)=793.1, *p*<0.001), with a similar increase in incorrect licking (high lick rate during the CS− cue period), during conditioning (Χ^2^(1)=9.89, *p*<0.01). **G)** Injections of AAV-packaged GFAP-Lck-GCaMP6f mixed with GFAP-cyto-tdTomato and GRIN lens implantation into the dmPFC (left) enabled visualization and *in vivo* recordings of individual astrocytic Ca^2+^ dynamics within an FOV (right). **H)** Examples of extracted Ca^2+^ traces from individual dmPFC astrocytes during a two-photon imaging session. **I, J)** Averaged traces from early (I) and late (J) conditioning show the population activity during CS+ and CS− trials across reward learning (Early: n=17 mice, 17 FOVs; 583 cells; Late: n=20 mice, 22 FOVs, 716 cells). **K, L)** Single-cell heatmaps from early **(K)** and late **(L)** behavioral sessions depict astrocytic recruitment and a refinement of cellular activity during CS+ trials throughout learning. **M)** Astrocytic activity discriminates between trial type, as single-cell auROC values significantly increased during CS+ trials, and simultaneously decreased during CS−trials, from early to late learning (Day x Trial Type interaction: F_1, 128597_= 9571.06, *p*<0.001). **N)** Tracked cells develop new excitatory responses during CS+ trials across learning, while responses in CS− trials decreased from early to late conditioning (Day x Trial Type interaction: F_1, 1064_=125.57, *p*<0.001). **O, P)** Direction-weighted distance correlations to the highest auROC values from CS+ and CS− trials across learning. In early learning, auROCs were not correlated with distance for either trial type **(O)**; however, by late learning auROCs from both CS+ and CS− trials negatively correlated with distance (P; Pearson-R value on each graph; Early: CS+ *p*=0.270, CS− *p*=0.083; Late: CS+ *p*<0.05, CS− *p*<0.001). **Q, R)** Correlation matrices of single-cell fluorescent signals within each trial type (CS+ or CS−) and subsequent inter-trial interval (ITI) in early **(Q)** and late **(R)** learning. There was greater response synchrony during CS+ trials that was strengthened across conditioning. **S)** Astrocytic responses during CS+ trials were highly correlated as compared to CS− trials, and CS+ response correlation significantly increased, while CS− response correlation decreased, across conditioning (Day x Trial Type interaction: F_1, 2594_=44.06, *p*<0.001). **T)** Astrocytic responses were most correlated in the ITIs following CS+ trials across learning, while response synchrony following CS− trials decreased with conditioning (Day x Trial Type interaction: F_1,2594_=17.036, *p*<0.001). **U)** During CS+ trials, the number of activated astrocytes significantly increased with learning (Fisher’s Exact Test, *p*<0.001), with no change in the number of significantly responding cells during CS− trials (Fisher’s Exact Test, *p*=0.105). **V)** Increased astrocytic Ca^2+^ responses reliably decode the trial type (CS+ or CS−), behavioral response (Correct Licking, Incorrect Withholding, Incorrect Licking, Correct Withholding), and reward delivery as compared to shuffled data during each behavioral session. Across learning, astrocytic decoding accuracy for CS+ and Correct Licking was maintained and significantly increased for Incorrect Withholding and cued reward delivery, while decoding accuracy decreased for CS−, Correct Withholding and Incorrect Licking (Day x Decoding Stimulus interaction: F_6,6990_=63.56, *p*<0.001). See also Figure S1. TI, trace interval; auROC, area under the receiver operating characteristic; FOV, field of view; ITI, inter-trial interval; Early, early Pavlovian conditioning (Day 1-2); Late, late Pavlovian conditioning (>0.65 licking auROC). Bar graphs are represented as mean ± SEM. Select statistical comparisons shown on graph, see Table 1 for complete statistical comparisons. Post-hoc comparisons: \**p*<0.05, \*\**p*<0.01, \*\*\**p*<0.001.

**Table 1 (Related to Figure 1).**
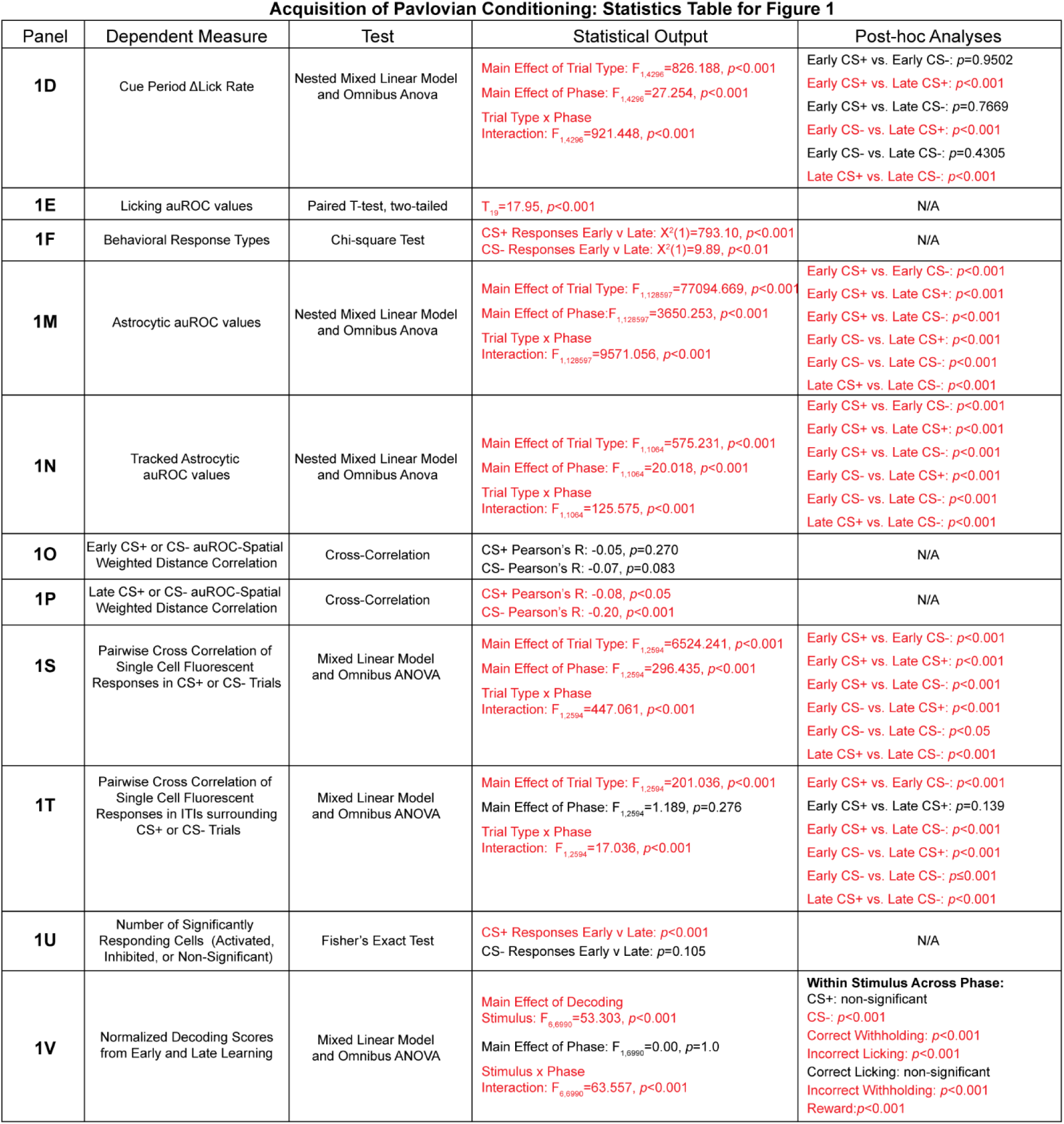
Statistical output for initial Pavlovian conditioning results presented in Figure 1.

The unique cellular morphology of astrocytes enables a single cell to directly contact, monitor, and regulate thousands of synapses ^36–39^, giving astrocytes the ability to modulate local neuronal activity as well as wider circuit-level dynamics. Considering this structural complexity and importance of Ca^2+^ transients in the peripheral processes for synaptic communication^40,41^, we employed an astrocyte-specific, membrane-targeted Ca^2+^ indicator (AAV5-GfaABC1D-Lck-GCaMP6f^27^) in combination with a cytosolic static label (AAV5-GfaABC1D-tdTomato) in our two-photon imaging experiments. The cytosolic tdTomato served as a static map of individual cell boundaries and was only captured during the pre- and post-behavioral session recordings (see methods). Thus, mice received injections of this viral cocktail into the dmPFC and implantation of a microendoscopic GRIN lens dorsal to the injection site, allowing visual access to GCaMP6f-expressing dmPFC astrocytes (**Fig. 1G, H; S1D**). Using two-photon microscopy, we measured acute changes in astrocytic GCaMP6f fluorescence during CS+ and CS− trials from early and late Pavlovian conditioning sessions. At both the population and single-cell level, dmPFC astrocytes displayed biased activation during CS+ trials, and these responses increased in magnitude and were further refined across reward learning (**Fig. 1I-L**). We then separated astrocytic fluorescent signals by behavioral response respective to each cue type. We found that GCaMP6f signal in early sessions was greatest during correct licking or incorrect withholding to CS+ presentation, followed next in magnitude by correct withholding to the CS−. Over the course of conditioning, the strength of fluorescent responses increased when mice correctly executed anticipatory licks or incorrectly withheld licking to the CS+, and responses dampened during correct withholding to the CS− (**Fig. S1 E-G**).

We then calculated the area under the receiver operating characteristic (auROC) curve for each cell based on trial-by-trial GCaMP6f fluorescence during behavioral imaging sessions. We compared fluorescence during a baseline period (3s) to that during an event window containing the CS, trace interval, and a post-sucrose period (**Fig. S1H**). These auROC values were then compared across trial types to evaluate evolving astrocytic discrimination between CS+ and CS− trials. In early learning, dmPFC astrocytic Ca^2+^ responses differentiated between trial types, showing stronger activation during CS+ trials. After conditioning, astrocytic activity continued to track the cue-sucrose association, with robust Ca^2+^ dynamics during CS+ trials that were accompanied by reduced responses specifically during CS− trials (**Fig. 1M**). These learning-related adaptations in Ca^2+^ responses were confirmed to occur within the same cells using auROC discrimination scores calculated from individual astrocytes orthogonally tracked from early to late Pavlovian conditioning sessions (**Fig. 1N, S1I**). To assess astrocytic Ca^2+^ dynamics after each trial, we assessed dmPFC astrocytic activity during equivalent time epochs within inter-trial intervals (ITIs) following CS+ and CS− trials. We found less Ca^2+^ activity during these ITIs and observed decreases in GCaMP6f fluorescence at the single-cell level (**Fig. S1J**). These results indicate that dmPFC astrocytes have acute Ca^2+^ transients that discriminate between trial types and dynamically track associative learning across conditioning.

Next, we quantified the relationship between spatial proximity and astrocytic Ca²⁺ response strength during CS⁺ and CS⁻ trials across early and late conditioning sessions. For each trial type, we ranked astrocytes by their auROC discrimination scores. We identified the cell with the strongest discrimination score and calculated distances from this reference cell to each cell within each FOV for each animal. These pairwise distances were then correlated with the corresponding auROC values for each astrocyte, whereby the strength of the negative correlation indicates spatial proximity of astrocytic discrimination within each trial type or corresponding ITI. We found that across conditioning, a significant distance-response relationship developed in both CS+ and CS− trials by late learning (**Fig. 1O, P**), indicating learning-dependent spatial refinement of astrocytic activity within each trial type. Although overall auROC discrimination was highest during CS+ trials, the spatial correlation was strongest in CS− trials during late learning, suggesting that weaker but more spatially organized Ca²⁺ responses developed for non-rewarded cues. We then examined direction-weighted spatial responses during the ITIs following each trial type. Despite the lower auROC values during ITIs as compared trials periods, we found significant distance-response relationships within the ITIs that followed each trial type (**Fig. S1K**). Considering this spatial refinement of astrocytic Ca^2+^ responses across learning, and that Ca^2+^ readily propagates through astrocytic networks via gap junctions^42^, we assessed the temporal synchrony between individual cells using pairwise cross-correlations during CS+ and CS− trials, as well as subsequent corresponding ITIs, across conditioning (**Fig. 1Q, R**). Astrocytic Ca^2+^ dynamics were most synchronized during CS+ trials, with network synchrony strengthened by cue-reward associative learning (**Fig. 1S**). During subsequent ITIs, responses were less synchronized than in the corresponding trials; however, early learning ITIs that followed CS+ trials had higher network synchrony than CS− trial ITIs, with response synchronization during CS+ ITIs strengthened with conditioning (**Fig. 1T**). To assess spatial relationships in response synchrony, we created 3-dimensional spatial maps wherein each cell was used as a central reference point, the corresponding cross-correlation values were heat mapped onto other cells using their individual ROI dimensions within each FOV. In this analysis, each cell contributed an individual layer and values from each pixel across each layer were averaged to create a pixel-by-pixel grid of spatially mapped cross correlation values. This was repeated for each FOV, for each trial type and ITI period (**Fig. S1L**). To generate a map depicting response similarity relative to cell proximity, we calculated the average cell diameter across the entire dataset and defined analysis zones based off this metric. To account for edge cases, the resulting correlation map was masked to 5-cell diameters, with pixels corresponding to cells outside of this spatial distance excluded from analysis. Distance-correlation data was calculated within each zone (the number of cell diameters from reference) during CS+ and CS− trials, as well as corresponding ITIs (**Fig. S1M-N**). We then isolated trial-specific relationships in response synchrony between astrocytes within each FOV. We calculated the spatially mapped average cross-correlation coefficient within each of 5-cell diameter zones for each trial type, corresponding subsequent ITIs, and the difference between the two for each cell (r_trial_-r_ITI_; **Fig. S1O-Q**). The difference between trial and subsequent ITI was then used to isolate trial-specific spatial response synchrony across learning. CS+ trials contained greater spatial response synchrony relative to CS− trials, and we observed learning-dependent recruitment of adjacent astrocytes throughout learning (**Fig. S1R**). Accordingly, we found that the number of activated cells during CS+ trials increased from early to late learning, while there was no change in CS− responding cells (**Fig. 1U**). Conversely, in the ITIs that followed CS+ trials, the number of astrocytes with inhibitory responses increased with conditioning, while there was no change in responding cells during the ITIs that followed CS− trials (**Fig. S1S**). Collectively, these data suggest that dmPFC astrocytes are recruited during associative learning, display new elevated Ca^+^ responses across conditioning, and develop coordinated, spatially organized Ca^2+^ patterns tuned toward the cue-reward pairing.

To assess whether dmPFC astrocytic Ca^2+^ dynamics encode specialized information pertaining to cues, reward, or CS−dependent behavioral responses, we used a machine learning-based SVM decoder to determine if astrocytic Ca^2+^ dynamics decoded task-relevant stimuli within each trial (**Fig. S1T**). We found decoder reliably predicted trial type (CS+ and CS−), behavioral response type (correct licking, incorrect withholding, correct withholding, incorrect licking), and reward delivery relative to chance during early and late conditioning sessions. Strikingly, in early sessions astrocytic responses had superior decoding ability for behavioral errors (incorrect licking to the CS−), which was refined downward with conditioning. Through conditioning, decoding accuracy remained constant for CS+ trials and correct licking responses, significantly increased for incorrect withholding and reward delivery, and decreased for CS− trials and CS− -dependent behavioral responses (**Fig. 1V**). These results indicate astrocytic responses shift from CS− related error detection to encoding CS+-induced behavioral responses and cued reward delivery. Together, our findings reveal that dmPFC astrocytes display time-locked activity patterns that track associative learning, develop new spatially coordinated Ca^2+^ dynamics as learning stabilizes, and encode correct and incorrect responses as behavior adapts across reward learning.

### dmPFC astrocytes preferentially encode the CS+-reward association

We found that dmPFC astrocytes display biased activation towards CS+ trials, but whether these responses are a result of CS+ presentation, reward delivery, or the combination of the two remained unclear. Thus, we ran well-trained mice through a series of omission tests: (1) sucrose omission test wherein there was a 50% probability of sucrose delivery following CS+ presentation or (2) cue omission that contained trials with un-cued sucrose delivery or no stimuli (**Fig. 2A**). Although there was a decrease in high-licking trials from late learning to the sucrose omission test (**Fig. S2A**), there was no change in trial discrimination (**Fig. 2B**) or CS+ correct rate (**Fig. S2B**) between these two sessions. Moreover, there was an increase in CS− correct rate from late Pavlovian conditioning to sucrose omission (**Fig. S2C**). Importantly, mice expressed conditioned licking behavior to CS+ presentation, regardless of reward status, that was followed by increased consummatory licks when reward was delivered (CS+ Rewarded) or lick cessation when sucrose was omitted (CS+ Unrewarded; **Fig. 2C, D**). These licking profiles were reflected in the behavioral response types during each trial (**Fig. 2E**), and indicated mice maintained anticipatory lick behavior despite 50% reward probability. During the cue omission test, mice only exhibited increased consummatory licks when reward was presented (**Fig. 2F, G**). These behavioral patterns enabled us to successfully assess the spatiotemporal dynamics of dmPFC across the variable trial types in the omission tests.

**Figure 2.**
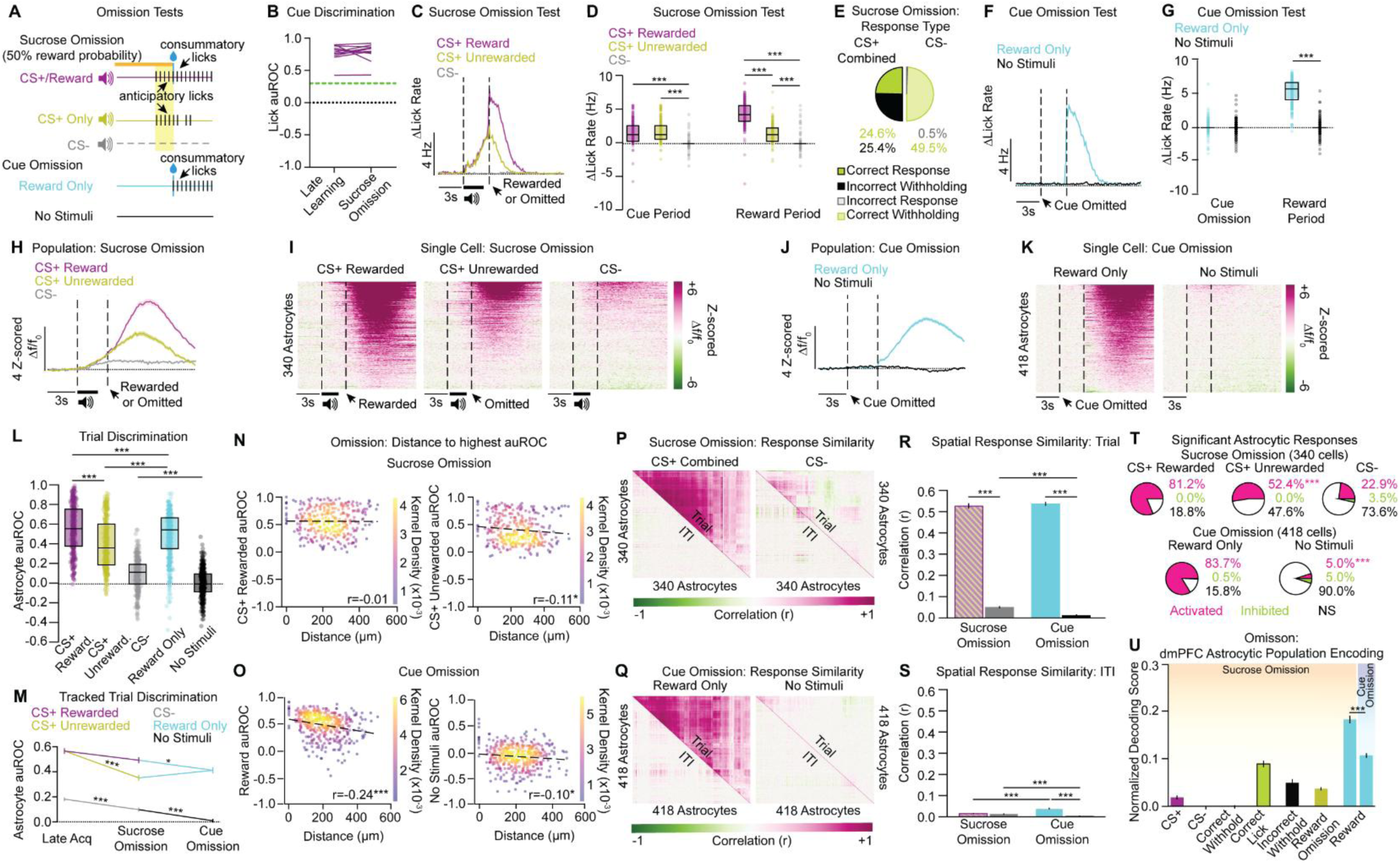
dmPFC astrocytes display the strongest activity during the cue-sucrose pairing, and encode cued sucrose delivery, as well as CS+-associated correct and incorrect behavioral responses. **A)** Behavioral schematic for the Omission Tests. During sucrose omission, there was a 50% probability that reward delivery would follow CS+ presentation, with no change to CS− trials. During cue omission, neither the CS+ or CS− was presented, and trials consisted of un-cued sucrose delivery or no stimuli. **B**) There was no change in behavioral trial discrimination from late learning to sucrose omission test (licking during the CS+ vs. CS− trace period; n=11 mice; T_10_=0.194, *p*=0.850). Green dashed line indicates an auROC threshold of 0.03. **C)** Averaged Δlick rate traces show that while there is no difference in anticipatory licking following CS+ presentation, the change in consummatory lick rate for CS+ Unrewarded trials drops off in comparison to CS+ Rewarded trials during the sucrose omission test (n=11 mice). **D)** There was no difference in the change in anticipatory lick rate during the cue period for CS+ Rewarded and Unrewarded trials, and both were significantly higher than CS− trials. During the reward period, the change in lick rate was the highest for CS+ Rewarded trials, followed by CS+ Unrewarded trials, then CS− trials (Period x Trial Type interaction: F_2,2184_=397.73, *p*<0.001). **E)** Sucrose omission test behavioral response types. **F)** Averaged Δlick rate traces show consummatory licks following un-cued sucrose delivery, and no licking when no stimuli were present (n=12 mice). **G)** As cue was omitted, there were no differences in the change in anticipatory lick rate during the cue period for rewarded and no stimuli trials, while during the reward period the change in lick rate increased following sucrose delivery relative to when there were no stimuli present (Period x Trial Type interaction: F_1,2385_=5159.99, *p*<0.001). **H**) Averaged fluorescent traces from sucrose omission test show the population activity during CS+ Rewarded, CS+ Unrewarded, and CS− trials (n=11 mice, 11 FOVs, 340 cells). **I)** Single-cell heatmaps from the sucrose omission test show the most robust cellular responses when sucrose delivery follows CS+ presentation, rather than when the CS+ is presented alone or during CS− trials. **J)** Averaged traces from the cue omission test show the population activity during un-cued sucrose delivery relative to when no stimuli are present. **K)** Single-cell heatmaps from the cue omission test depict strong cellular activation following un-cued sucrose delivery and few Ca^2+^ dynamics when no stimuli are present. **L)** Astrocytic auROC values from each trial type during the omission tests. Astrocytic discrimination was highest in trials where the CS+ was paired with sucrose delivery, followed next by Reward Only trials, then CS+ Unrewarded trials, then CS− trials, and finally by trials where no stimuli were present (Nested One-way ANOVA: F_4,1851_=512.43, *p*<0.001). **M)** Tracked astrocytic auROC values from late Pavlovian conditioning to the omission tests. Tracked astrocytes respond most strongly when CS+ predicts reward delivery. There were no differences in trial discrimination from late learning to sucrose omission CS+ Rewarded trials, and these were both significantly higher than trials where CS+ and reward were presented alone. For CS− trials, trial discrimination was highest in late acquisition, followed by CS− trials in sucrose omission, and then trials when there were no stimuli present (Nested One-way ANOVA: F_6,903_=114.66, *p*<0.001). **N)** During the sucrose omission test, when CS+ preceded sucrose delivery there was no correlation between auROC and distance (Pearson-R on graph, *p*=0.910), while there was a negative auROC-distance correlation when CS+ was presented alone (Pearson-R on graph, *p*<0.05). **O)** In the cue omission test, there was a significant auROC-distance correlation when reward was presented alone (Pearson-R on graph, *p*<0.001) as well as when no stimuli were present (Pearson-R on graph, *p*<0.05). **P)** Correlation matrices of single-cell fluorescent signals within CS+ (combined across rewarded and unrewarded trials) and CS− trials, as well as subsequent ITIs, during sucrose omission show greatest response similarity during CS+ trials combined. **Q)** Correlation matrices of single-cell fluorescent signals within Reward Only and No Stimuli trials, as well as subsequent ITIs, depict greatest response similarity during Reward Only trials. **R)** Astrocytic responses from sucrose omission CS+ trials combined and cue omission Reward Only trials were highly correlated as compared to CS− or No Stimuli trials, with no differences between CS+ combined or Reward Only trials. CS− response correlation was higher than trials with no stimuli (Day x Trial Type Interaction: F_1,1512_=16.67, *p*<0.001). **S)** Astrocytic responses were most correlated in the ITIs following Reward Only trials, followed next by ITIs that were preceded by CS+ trials (combined), then by CS− trials (Day x Trial Type Interaction: F_1,1512_=147.278, *p*<0.001). **T)** There were significantly less activated cells when CS+ was presented alone than during CS+ Rewarded or Reward Only trials (CS+ Trials Fisher’s Exact test *p*<0.001). CS− trials from sucrose omission had significantly more activated cells than when no stimuli were present (CS− Trials Fisher’s Exact test, *p*<0.001). **U)** During sucrose omission, astrocyte activation accurately decoded CS+ trial type, correct licking and incorrect withholding, and reward omission or delivery as compared to shuffled data. Increased astrocytic Ca^2+^ also reliably decoded reward delivery during the cue omission test, but with less accuracy than CS+-predicted reward delivery in sucrose omission (Day x Decoding Stimulus interaction: F_6,4317_=22.55, *p*<0.001). See also Figure S2. TI, trace interval; auROC, area under the receiver operating characteristic; ITI, inter-trial interval. Bar graphs are represented as mean ± SEM. Select statistical comparisons shown on graph, see Table 2 for complete statistical comparisons. Post-hoc comparisons: \**p*<0.05, \*\**p*<0.01, \*\*\**p*<0.001.

**Table 2 (Related to Figure 2).** Statistical output for Omission Test results presented in Figure 2 (Omission Tests).

| Panel | Dependent Measure | Test | Statistical Output | Post-hoc Analyses |
| --- | --- | --- | --- | --- |
| **2B** | Licking auROC values | Paired T-test, two-tailed | $T_{11}=0.194, p=0.8499$ | N/A |
| **2D** | Sucrose Omission:<br>Cue Period ΔLick Rate | Nested Mixed Linear Model and Omnibus Anova | Main Effect of Trial Type: $F_{2,2184}=1557.350, p<0.001$<br>Main Effect of Period: $F_{1,2184}=209.816, p<0.001$<br>Trial Type x Period Interaction: $F_{2,2184}=397.731, p<0.001$ | Cue CS+ Rewarded vs. Cue CS+ Unrewarded: $p=0.513$<br>Cue CS+ Rewarded vs. Cue CS-: $p<0.001$<br>Cue CS+ Rewarded vs. Reward CS+ Rewarded: $p<0.001$<br>Cue CS+ Rewarded vs. Reward CS+ Unrewarded: $p=0.290$<br>Cue CS+ Rewarded vs. Reward CS-: $p<0.001$<br>Cue CS+ Unrewarded vs. Cue CS-: $p<0.001$<br>Cue CS+ Unrewarded vs. Reward CS+ Rewarded: $p<0.001$<br>Cue CS+ Unrewarded vs. Reward CS+ Unrewarded: $p<0.01$<br>Cue CS+ Unrewarded vs. Reward CS-: $p<0.001$<br>Cue CS- vs. Reward CS+ Rewarded: $p<0.001$<br>Cue CS- vs. Reward CS+ Unrewarded: $p<0.001$<br>Cue CS- vs. Reward CS-: $p=0.995$<br>Reward CS+ Rewarded vs. Reward CS+ Unrewarded: $p<0.001$<br>Reward CS+ Rewarded vs. Reward CS-: $p<0.001$<br>Reward CS+ Unrewarded vs. Reward CS-: $p<0.001$ |
| **2G** | Cue Omission:<br>Cue Period ΔLick Rate | Nested Mixed Linear Model and Omnibus Anova | Main Effect of Trial Type: $F_{1,2385}=5242.514, p<0.001$<br>Main Effect of Period: $F_{1,2385}=5123.296, p<0.001$<br>Trial Type x Period Interaction: $F_{1,2385}=5159.999, p<0.001$ | Cue Omit Reward vs. Cue Omit No Stimuli: $p=0.9801$<br>Cue Omit Reward vs. Reward Reward: $p<0.001$<br>Cue Omit Reward vs. Reward No Stimuli: $p=0.9433$<br>Cue Omit No Stimuli vs. Reward Reward: $p<0.001$<br>Cue Omit No Stimuli vs. Reward No Stimuli: $p=0.9981$<br>Reward Reward vs. Reward No Stimuli: $p<0.001$ |
| **2L** | Astrocytic auROC values | Nested Mixed Linear Model and Omnibus One-way Anova | Main Effect of Trial Type: $F_{4,1855}=512.431, p<0.001$ | CS+ Unrewarded vs. CS+ Rewarded: $p<0.001$<br>CS+ Unrewarded vs. CS-: $p<0.001$<br>CS+ Unrewarded vs. Reward Only: $p<0.001$<br>CS+ Unrewarded vs. No Stimuli: $p<0.001$<br>CS+ Rewarded vs. CS-: $p<0.001$<br>CS+ Rewarded vs. Reward Only: $p<0.001$<br>CS+ Rewarded vs. No Stimuli: $p<0.001$<br>CS- vs. Reward Only: $p<0.001$<br>CS- vs. No Stimuli: $p<0.001$<br>Reward Only vs. No Stimuli: $p<0.001$ |
| **2M** | Tracked Astrocytic auROC values | Nested Mixed Linear Model and Omnibus One-way Anova | Main Effect of Trial Type: $F_{6,903}=114.675, p<0.001$ | Late CS+ vs. Late CS-: $p<0.001$<br>Late CS+ vs. CS+ Unrewarded: $p<0.001$<br>Late CS+ vs. CS+ Rewarded: $p=0.1097$<br>Late CS+ vs. Suc Omit CS-: $p<0.001$<br>Late CS+ vs. Reward Only: $p<0.001$<br>Late CS+ vs. No Stimuli: $p<0.001$<br>Late CS- vs. CS+ Unrewarded: $p<0.001$<br>Late CS- vs. CS+ Rewarded: $p<0.001$<br>Late CS- vs. Suc Omit CS-: $p<0.05$<br>Late CS- vs. Reward Only: $p<0.001$<br>Late CS- vs. No Stimuli: $p<0.001$<br>CS+ Unrewarded vs. CS+ Rewarded: $p<0.001$<br>CS+ Unrewarded vs. Suc Omit CS-: $p<0.001$<br>CS+ Unrewarded vs. Cue Only: $p=0.3055$<br>CS+ Unrewarded vs. No Stimuli: $p<0.001$<br>CS+ Rewarded vs. Suc Omit CS-: $p<0.001$<br>CS+ Rewarded vs. Reward Only: $p<0.05$<br>CS+ Rewarded vs. No Stimuli: $p<0.001$<br>Suc Omit CS- vs. Reward Only: $p<0.001$<br>Suc Omit CS- vs. No Stimuli: $p<0.05$<br>Reward Only vs. No Stimuli: $p<0.001$ |
| **2N** | Sucrose Omission auROC-Spatial Weighted Distance Correlation | Cross-Correlation | CS+ Rewarded Pearson's R: $-0.01, p=0.910$<br>CS+ Unrewarded Pearson's R: $-0.11, p<0.05$<br>CS- Pearson's R: $-0.15, p<0.01$ | N/A |
| **2O** | Cue Omission auROC-Spatial Weighted Distance Correlation | Cross-Correlation | Reward Only Pearson's R: $-0.24, p<0.001$<br>No Stimuli Pearson's R: $-0.10, p<0.05$ | N/A |
| **2R** | Pairwise Cross Correlation of Single Cell Fluorescent Responses Across the Omission Test Trial Types | Mixed Linear Model and Omnibus ANOVA | Main Effect of Day: $F_{1,1512}=5.692, p<0.05$<br>Main Effect of Trial Type: $F_{1,1512}=7261.136, p<0.001$<br>Trial Type x Day Interaction: $F_{1,1512}=16.667, p<0.001$ | Suc Omit CS+ vs. Suc Omit CS-: $p<0.001$<br>Suc Omit CS+ vs. Reward Only: $p=0.6271$<br>Suc Omit CS+ vs. No Stimuli: $p<0.001$<br>Suc Omit CS- vs. Reward Only: $p<0.001$<br>Suc Omit CS- vs. No Stimuli: $p<0.001$<br>Reward Only vs. No Stimuli: $p<0.001$ |
| **2S** | Pairwise Cross Correlation of Single Cell Fluorescent Responses Across the Omission Test ITIs | Mixed Linear Model and Omnibus ANOVA | Main Effect of Day: $F_{1,1512}=28.398, p<0.001$<br>Main Effect of Trial Type: $F_{1,1512}=238.225, p<0.001$<br>Trial Type x Day Interaction: $F_{1,1512}=147.278, p<0.001$ | Suc Omit CS+ ITI vs. Suc Omit CS- ITI: $p=0.547$<br>Suc Omit CS+ ITI vs. Reward Only ITI: $p<0.001$<br>Suc Omit CS+ ITI vs. No Stimuli ITI: $p<0.001$<br>Suc Omit CS- ITI vs. Reward Only ITI: $p<0.001$<br>Suc Omit CS- ITI vs. No Stimuli ITI: $p<0.001$<br>Reward Only ITI vs. No Stimuli ITI: $p<0.001$ |
| **2T** | Number of Significantly Responding Cells (Activated, Inhibited, or Non-Significant) | Fisher's Exact Test | CS+ Rewarded, CS+ Unrewarded, Reward Only Responses: $p<0.001$<br>CS- vs. No Stimuli Responses: $p<0.001$ | N/A |
| **2U** | Normalized Decoding Scores from the Omission Tests | Mixed Linear Model and Omnibus ANOVA | Main Effect of Decoding Stimulus: $F_{6,4317}=242.588, p<0.001$<br>Main Effect of Phase: $F_{1,4317}=0.00, p=1.0$<br>Stimulus x Phase Interaction: $F_{6,4317}=22.546, p<0.001$ | **Relevant Within Day Across Stimuli:**<br>CS+-predicted reward vs. reward omission: $p<0.001$<br>**Relevant Within Stimuli Across Day:**<br>CS+-predicted reward vs. Reward Only: $p<0.001$ |

We used two-photon Ca^2+^ imaging to examine acute astrocytic responses during CS+ Rewarded, CS+ Unrewarded, CS−, Reward Only, and No Stimuli trials. At both the population and single-cell level, dmPFC astrocytes display robust Ca^2+^ elevations during trials when CS+ precedes sucrose delivery as opposed to when CS+ or sucrose are presented alone (**Fig. 2H-K**), regardless of CS+-associated behavioral response (correct licking, incorrect withholding; **Fig. S2D, E**). Accordingly, astrocytic response strength was greatest during the CS+-sucrose pairing, followed next in magnitude by Reward Only trials, and then CS+ Unrewarded trials (**Fig. 2L**), indicating greatest astrocytic discrimination for CS+ Rewarded trials. To assess how individual cells responded across each trial type, we longitudinally tracked astrocytic auROC values from late Pavlovian conditioning through the omission tests. While there was no change in trial discrimination between cells tracked from late CS+ trials to CS+ Rewarded trials, cells decreased responding during CS+ or sucrose only trials (**Fig. 2M; S2F, G**). Astrocytic responses during all stimulus-containing trials exceeded subsequent ITI activity (**Fig. S2H**). We then examined the spatial relationship of astrocytic response strength within the omission tests. We found that when either CS+ or reward was presented alone, there was a significant auROC response-distance relationship that was not present in CS+ Rewarded trials, indicating when reward is unpredictable there is a spatial refinement of dmPFC astrocytic activity. We also observed response strength-distance correlations during no stimuli trials in cue omission, as well as during the ITIs following the variable trial types during the omission tests (**Fig. 2N, O; S2I**), implicating dmPFC astrocytes in PFC feedback monitoring during the time periods that surround stimuli-containing events.

We next determined temporal synchrony in the omission tests. Given mice were exposed to the reward-paired stimuli with no differences in conditioned lick behavior during the sucrose omission test, fluorescent signals were combined for CS+ temporal and spatial synchrony analyses regardless of reward status (rewarded or omitted). Using individual cell-pairwise cross-correlations, we found that there is high astrocytic response synchrony during CS+ combined and Reward Only trials relative to their subsequent respective ITI periods (**Fig. 2P, Q**), with no differences in correlation strength between these trial types (**Fig. 2R**). Although there was overall less response synchrony during ITI periods, the ITIs following Reward Only trials had the greatest astrocytic Ca^2+^ event synchrony relative to ITIs that were preceded by CS+ combined or CS− trials (**Fig. 2S**), suggesting coordinated astrocytic network activity in ITIs following unpredictable sucrose delivery. We then mapped the average cross-correlations from each trial type and subsequent ITI spatially to assess response synchrony within adjacent astrocytes (**Fig. S2J, K**). We isolated trial-specific response synchrony by determining the difference in average correlation coefficient for each cell during CS+ combined, CS−, Reward Only, and No Stimuli trials relative to their respective ITI periods (**Fig. S2L**). There were no differences in network synchronization between CS+ combined trials and Reward Only trials, with both having greater spatial network synchrony than CS− or No Stimuli trials (**Fig. S2M**). These data suggest that there are strongly coordinated Ca^2+^ responses throughout the astrocytic syncytia when CS+, sucrose, or both are present.

Next, we examined the number of significantly responding astrocytes and the decoding accuracy of these cells across the omission tests. We found that CS+ Rewarded and Reward Only trials recruited more activated astrocytes than CS+ presentation alone. Moreover, there were more activated cells during CS− trials than when no stimuli were present (**Fig. 2T**). During the subsequent ITIs, a greater number of astrocytes were significantly inhibited in the ITIs following Reward Only trials as compared to the ITIs preceded by CS+ combined trials. Further, there were significantly more activated astrocytes in the ITIs that followed No Stimuli trials as compared to CS− -linked ITIs (**Fig. S2N**). Collectively, these data demonstrate that astrocytic responses are strongest and time-locked to stimulus-containing trials, rather than the subsequent ITI periods. Using our machine learning-based decoder, we determined that astrocytic responses accurately decode CS+ trials, CS+-associated behavioral responses (correct lick, incorrect withholding), and reward delivery and omission compared to chance. Moreover, astrocytes had superior decoding accuracy for reward that was preceded by CS+ relative to un-cued sucrose delivery (**Fig. 2U**). Collectively, these results establish dmPFC astrocytes preferentially encode the cue-reward association.

### dmPFC astrocytes rapidly track updated reward contingencies and encode new cue-reward associations, as well as updated and outdated motivated behavioral action, after reversal learning

Based on our findings that dmPFC astrocytic Ca^2+^ responses track associative learning and preferentially encode the cue-reward association and conditioned behavior, we predicted that astrocytic dynamics would shift to reflect reversal of the cue-reward pairing and subsequent behavioral adaptation. Thus, we used two-photon Ca^2+^ imaging to measure astrocytic activity patterns during head-fixed Pavlovian conditioning sessions, wherein well-trained mice learned to associate the old CS− with sucrose delivery (new CS+) and the old CS+ was no longer paired with reward (**Fig. 3A**). With reversal training, mice appropriately adapted behavior and extinguished conditioned licking to the old CS+, while increasing anticipatory licking following presentation of the new CS+ by late learning (**Fig. 3B, C; S3A-C**). This was reflected in the strong trial discrimination in late reversal (**Fig. 3D**), where behavioral responses had shifted from old CS+ licking errors to correct responses following the new sucrose-paired cue (**Fig. 3E**). Taken together, these results demonstrate mice learned the updated cue-sucrose association and appropriately adapted behavior in response to both the old and new, CS+.

**Figure 3.**
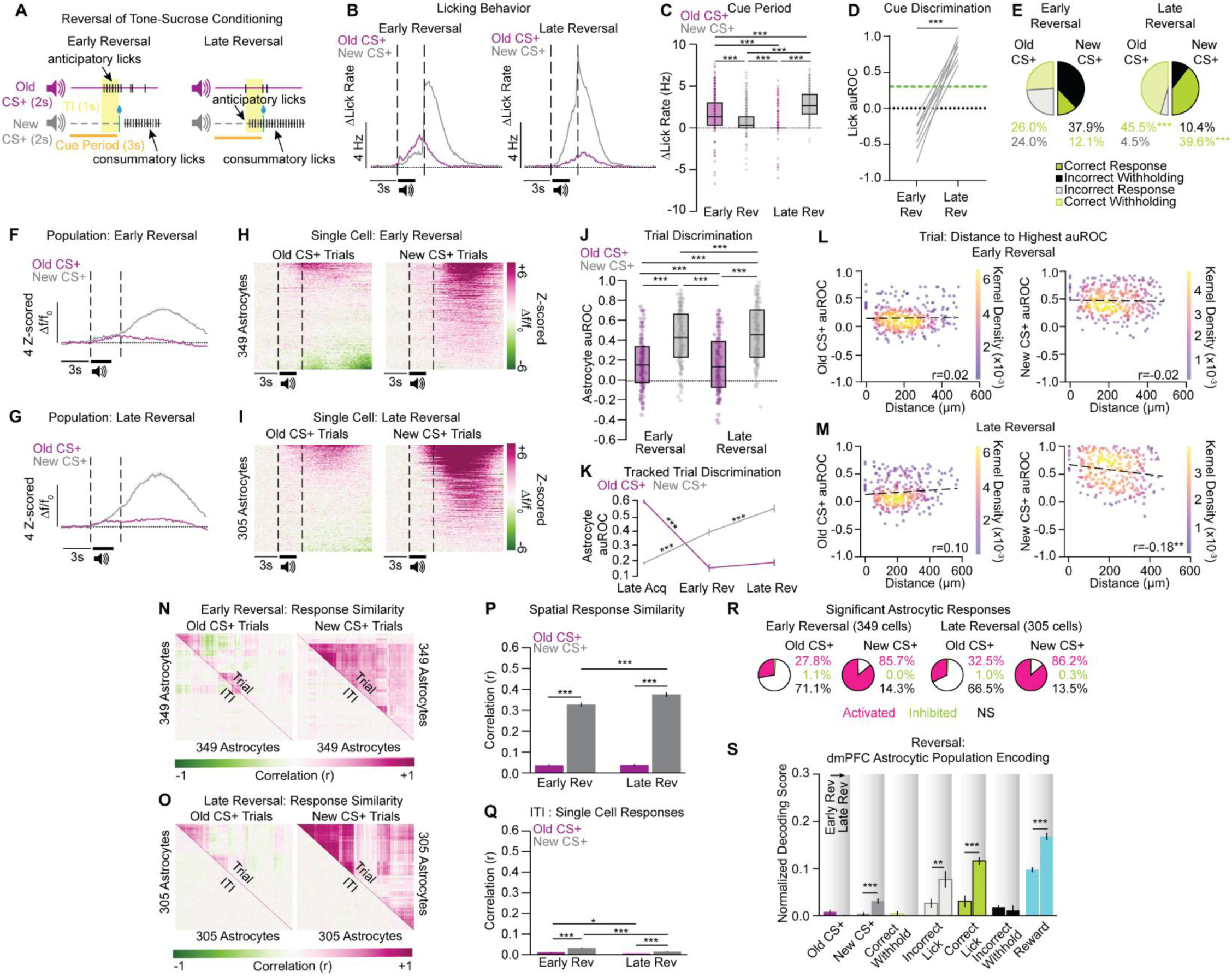
dmPFC astrocytic activity rapidly adapts to track new CS+-sucrose association and encodes the new cue-reward pairing and updated, as well as outdated, motivated behavioral action. **A)** Behavioral schematic for reward contingency updating and reversal learning. **B)** Averaged Δlick rate traces show that anticipatory licking shifts from the old CS+ in early (left) to the new CS+ across in late (right) reversal learning (n=8-9 mice). **C)** The change in anticipatory lick rate during the cue period was significantly higher following presentation of the old CS+ relative to the new CS+ in early reversal but adapted across conditioning to be significantly elevated following presentation of the new CS+ (n=8-9 mice; Day x Trial Type interaction: F_1,1896_=610.08, *p*<0.001). **D)** Behavioral discrimination (anticipatory licking to the new CS+ vs old CS+) significantly increased from early to late reversal (T_7_=14.88, *p*<0.001). Green line indicates auROC threshold of 0.3. **E)** Correct withholding, low licking during the old CS+ cue period, and correct licking, anticipatory licks during the new CS+ cue period, both significantly increased from early to late reversal learning (Correct withholding: Χ ^2^(1)=155.44, *p*<0.001; Correct licking: Χ^2^ (1)=279.77, *p*<0.001). **F, G)** Averaged traces from early (**F**) and late (**G**) reversal conditioning show the population activity during old and new CS+ trials as mice learn the updated reward contingency (Early Rev: n=9 mice, 10 FOVs, 349 cells; Late Rev: n=8 mice, 9 FOVs, 305 cells). **H, I)** Single-cell heatmaps from early (**H**) and late (**I**) reversal sessions depict rapid Ca^2+^ response adaptations during new CS+ trials that is refined across learning. **J)** Single-cell activity discriminates between old and new CS+ trial types, as astrocytic auROC values are significantly greater during new CS+ compared to old CS+ trials. Across reversal learning, astrocytic response strength increases during new, and decreases during old, CS+ trials (Day x Trial Type interaction: F_1, 64742_= 494.06, *p*<0.001). **K)** Astrocytes tracked from late Pavlovian conditioning through reversal learning. When the CS− became the new CS+, astrocytes rapidly developed new activated responses, and these strengthened across reversal training. When the CS+ no longer predicted reward delivery, there was a dramatic decrease in astrocytic response strength during old CS+ trials relative to Late Acquisition (Day X Trial Type interaction: F_2,366_=115.131, *p*<0.001). **L, M)** Correlations of direction-weighted distance with auROC values from old and new CS+ trials across reversal learning. In early reversal, there was no distance-auROC relationship during either the old or new CS+ trials (**L**; Pearson-R value on each graph; old CS+ *p*=0.75; new CS+ *p*=0.65). Following reversal conditioning, there was a significant distance-auROC correlation during new, but not old, CS+ trials (**M**; Pearson-R value on each graph; old CS+ *p*=0.091; new CS+ *p*<0.01). **N, O)** Correlation matrices of single-cell fluorescent signals within each trial type (old or new CS+) and subsequent inter-trial intervals (ITIs) in early (**N**) and late (**O**) reversal depict greater response similarity during new CS+ trials that strengthens with conditioning. **P)** Astrocytic fluorescent responses were highly correlated during new CS+ trials relative to old CS+ trials, with response correlations increasing during new CS+ trials with reversal training (Day x Trial Type interaction: F_1, 1304_=15.52, *p*<0.001). **Q)** Astrocytic responses were most correlated in the ITIs following new CS+ trials in early reversal, with these correlations decreased by late reversal. Response similarity decreased in ITIs following the old CS+ across reversal learning (Day x Trial Type interaction: F_1,1304_=25.014, *p*<0.001). **R)** There was no change in the number of significantly responding astrocytes during old CS+ or new CS+ trials from early to late reversal (old CS+: Fisher’s Exact Test, *p*=0.418; new CS+: Fisher’s Exact Test, *p*=0.648). **S)** Increased astrocytic Ca^2+^ reliably decoded old CS+ trials, incorrect behavioral responses (Incorrect Licking to old CS+, incorrect withholding to new CS+), and reward delivery as compared to chance in early reversal. After mice learn the updated reward contingency, astrocytic decoding accuracy increased for new CS+ trials, correct and incorrect behavioral responses (Correct Licking to new CS+, Incorrect Licking to old CS+), and cued reward delivery relative to early reversal scores (Day x Decoding Stimulus interaction: F_6,3706_=21.50, *p*<0.001). See also Figure S3. TI, trace interval; auROC, area under the receiver operating characteristic; ITI, inter-trial interval; Early Rev, early reversal learning (Reversal Day 1-2); Late Rev, late reversal learning (>0.65 licking auROC). Bar graphs are represented as mean ± SEM. Select statistical comparisons shown on graph, see Table 3 for complete statistical comparisons. Post-hoc comparisons: \**p*<0.05, \*\**p*<0.01, \*\*\**p*<0.001.

**Table 3 (Related to Figure 3).** Statistical output for Reversal Learning results presented in Figure 3.

| Panel | Dependent Measure | Test | Statistical Output | Post-hoc Analyses |
| --- | --- | --- | --- | --- |
| 3C | Cue Period ΔLick Rate | Nested Mixed Linear Model and Omnibus Anova | Main Effect of Phase: $F_{1,1898}=0.236, p=0.627$<br>Main Effect of Trial Type: $F_{1,1898}=97.471, p<0.001$<br>Phase x Trial Type<br>Interaction: $F_{1,1898}=610.076, p<0.001$ | Rev-Early old CS+ vs. Rev-Early new CS+: $p<0.001$<br>Rev-Early old CS+ vs. Rev-Late old CS+: $p<0.001$<br>Rev-Early old CS+ vs. Rev-Late new CS+: $p<0.001$<br>Rev-Early new CS+ vs. Rev-Late old CS+: $p<0.001$<br>Rev-Early new CS+ vs. Rev-Late new CS+: $p<0.001$<br>Rev-Late old CS+ vs. Rev-Late new CS+: $p<0.001$ |
| 3D | Licking auROC values | Paired T-test, two-tailed | $T_7=14.88, p<0.001$ | N/A |
| 3E | Behavioral Response Types | Chi-square Test | Old CS+ Responses Rev-Early vs. Rev-Late: $\chi^2(1)=155.44, p<0.001$<br>New CS+ Responses Rev-Early vs. Rev-Late: $\chi^2(1)=279.77, p<0.001$ | N/A |
| 3J | Astrocytic auROC values | Nested Mixed Linear Model and Omnibus Anova | Main Effect of Phase: $F_{1,64742}=1081.464, p<0.001$<br>Main Effect of Trial Type: $F_{1,64742}=36116.238, p<0.001$<br>Phase x Trial Type<br>Interaction: $F_{1,64742}=494.064, p<0.001$ | Rev-Early old CS+ vs. Rev-Early new CS+: $p<0.001$<br>Rev-Early old CS+ vs. Rev-Late old CS+: $p<0.001$<br>Rev-Early old CS+ vs. Rev-Late new CS+: $p<0.001$<br>Rev-Early new CS+ vs. Rev-Late old CS+: $p<0.001$<br>Rev-Early new CS+ vs. Rev-Late new CS+: $p<0.001$<br>Rev-Late old CS+ vs. Rev-Late new CS+: $p<0.001$ |
| 3K | Tracked Astrocytic auROC values from Late Acquisition through Reversal | Nested Mixed Linear Model and Omnibus Anova | Main Effect of Phase: $F_{2,398}=14.538, p<0.001$<br>Main Effect of Trial Type: $F_{1,398}=9.364, p<0.01$<br>Phase x Trial Type<br>Interaction: $F_{2,398}=115.131, p<0.001$ | Late CS+ vs. Late CS-: $p<0.001$<br>Late CS+ vs. Rev-Early old CS+: $p<0.001$<br>Late CS+ vs. Rev-Early new CS+: $p<0.001$<br>Late CS+ vs. Rev-Late old CS+: $p<0.001$<br>Late CS+ vs. Rev-Late new CS+: $p=0.541$<br>Late CS- vs. Rev-Early Old CS+: $p=0.578$<br>Late CS- vs. Rev-Early new CS+: $p<0.001$<br>Late CS- vs. Rev-Late old CS+: $p=0.981$<br>Late CS- vs. Rev-Late new CS+: $p<0.001$<br>Rev-Early old CS+ vs. Rev-Early new CS+: $p<0.001$<br>Rev-Early old CS+ vs. Rev-Late old CS+: $p=0.941$<br>Rev-Early old CS+ vs. Rev-Late new CS+: $p<0.001$<br>Rev-Early old CS+ vs. Rev-Late old CS+: $p<0.001$<br>Rev-Early new CS+ vs. Rev-Late new CS+: $p<0.001$<br>Rev-Late old CS+ vs. Rev-Late new CS+: $p<0.001$ |
| 3L | Early Reversal old and new CS+ auROC-Spatial Weighted Distance Cross-Correlation | Cross-Correlation | old CS+ Pearson's R: 0.02, $p=0.750$<br>new CS+ Pearson's R: -0.08, $p=0.160$ | N/A |
| 3M | Late Reversal old and new CS+ auROC-Spatial Weighted Distance Cross-Correlation | Cross-Correlation | old CS+ Pearson's R: 0.10, $p=0.091$<br>new CS+ Pearson's R: -0.18, $p<0.01$ | N/A |
| 3P | Pairwise Cross Correlation of Single Cell Fluorescent Responses in old and new CS+ Trials | Mixed Linear Model and Omnibus ANOVA | Main Effect of Phase: $F_{1,1304}=16.45, p<0.001$<br>Main Effect of Trial Type: $F_{1,1304}=2706.222, p<0.001$<br>Phase x Trial Type<br>Interaction: $F_{1,1304}=15.520, p<0.001$ | Rev-Early old CS+ vs. Rev-Early new CS+: $p<0.001$<br>Rev-Early old CS+ vs. Rev-Late old CS+: $p=0.999$<br>Rev-Early old CS+ vs. Rev-Late new CS+: $p<0.001$<br>Rev-Early new CS+ vs. Rev-Late old CS+: $p<0.001$<br>Rev-Early new CS+ vs. Rev-Late new CS+: $p<0.001$<br>Rev-Late old CS+ vs. Rev-Late new CS+: $p=0.578$ |
| 3O | Pairwise Cross Correlation of Single Cell Fluorescent Responses in ITIs surrounding old and new CS+ Trials | Mixed Linear Model and Omnibus ANOVA | Main Effect of Phase: $F_{1,1304}=87.297, p<0.001$<br>Main Effect of Trial Type: $F_{1,1304}=156.197, p<0.001$<br>Phase x Trial Type<br>Interaction: $F_{1,1304}=25.014, p<0.001$ | Rev-Early old CS+ ITI vs. Rev-Early new CS+ ITI: $p<0.001$<br>Rev-Early old CS+ ITI vs. Rev-Late old CS+ ITI: $p<0.05$<br>Rev-Early old CS+ ITI vs. Rev-Late new CS+ ITI: $p=0.199$<br>Rev-Early new CS+ ITI vs. Rev-Late old CS+ ITI: $p<0.01$<br>Rev-Early new CS+ ITI vs. Rev-Late new CS+ ITI: $p<0.001$<br>Rev-Late old CS+ ITI vs. Rev-Late new CS+ ITI: $p<0.01$ |
| 3R | Number of Significantly Responding Cells (Activated, Inhibited, or Non-Significant) | Fisher's Exact Test | old CS+ Responses Rev-Early v Rev-Late: $p=0.418$<br>new CS+ Responses Rev-Early v Rev-Late: $p=0.648$ | N/A |
| 3S | Normalized Decoding Scores from Early and Late Learning | Mixed Linear Model and Omnibus ANOVA | Main Effect of Decoding<br>Stimulus: $F_{8,3708}=34.982, p<0.001$<br>Main Effect of Phase: $F_{1,3708}=0.00, p=1.0$<br>Stimulus x Phase<br>Interaction: $F_{8,3708}=21.501, p<0.001$ | Within Stimulus Across Phase:<br>old CS+: non-significant<br>new CS+: $p<0.001$<br>Correct Withholding: non-significant<br>Incorrect Licking: $p<0.01$<br>Correct Licking: $p<0.001$<br>Incorrect Withholding: non-significant<br>Reward: $p<0.001$ |

Given behavior appropriately updated to reflect the new cue-sucrose association, we used two-photon microscopy to examine acute dmPFC astrocytic Ca^2+^ dynamics during old and new CS+ trials across reversal learning. At both the population and single-cell level, astrocytic responses immediately shifted toward the new CS+-sucrose pairing, and this biased activation strengthened with conditioning (**Fig. 3F-I**). Upon reward contingency reversal, GCaMP6f fluorescence was strongest in new CS+ trials when mice incorrectly withheld responding; however, by late reversal Ca^2+^ responses were greatest when mice appropriately licked to the new CS+. Moreover, there was a learning-dependent dampening of Ca^2+^ activity during trials when mice displayed incorrect anticipatory licking to the old CS+ (**Fig. S3D-F**), indicating decreased astrocytic activity as the old CS+ lost incentive salience. These results are consistent with our initial findings and support the idea that astrocytes function as error detection units during early associative learning that adapt cellular responses to reflect appropriate behavioral updating. Accordingly, dmPFC astrocytic strength increased during new CS+ trials as mice learned the updated contingency (**Fig. 3J**), indicating astrocytic activity reliably discriminates between trial types. We tracked individual cells from late Pavlovian conditioning through reversal learning, and found astrocytes immediately developed new, activated responses to the new CS+-sucrose pairing that strengthened with conditioning. Strikingly, in these same cells, there was a dramatic decrease in Ca^2+^ activity during old CS+ trials when tracked from late acquisition (**Fig. 3K; S3G**). Further, population activity was time-locked to trials (**Fig. S3H**).

We then quantified the relationship between spatial distance and astrocytic response strength across reversal learning. We found there was a direction-weighted spatial response relationship selectively present in new CS+ trials during late reversal (**Fig. 3L, M**), indicating learning-dependent spatial refinement of Ca^2+^ dynamics within the astrocytic syncytia. In early reversal, there was a correlation between astrocytic response strength and spatial distance present in the trials that followed new, but not old, CS+ trials, suggesting spatially refined astrocytic Ca^2+^ signals in ITIs that followed trials with the updated cue-reward pairing. By late reversal, ITIs that followed both old and new CS+ trials had significant spatial distance-response relationships (**Fig. S3I**). Considering the strong directional component in response strength during trials and subsequent ITIs, we examined response synchrony across individual cells using pairwise cross-correlations during old and new CS+ trials across reversal (**Fig. 3N, O**). Astrocytic Ca^2+^ signals were highly synchronized during new CS+ trials, with this synchronization strengthening across reversal learning (**Fig. 3P**). Consistently, we found response synchrony was highest during ITIs that followed new CS+ trials in early reversal that was refined down with training (**Fig. 3Q**). These data indicate greatest astrocytic response synchrony during trials with the new CS+-sucrose pairing, as well as in the ITIs that follow. Next, we constructed spatial maps of the average cross-correlation coefficients from each trial type, subsequent corresponding ITI, and the difference between the two to determine trial-specific spatial response synchronization across reversal learning (**Fig. S3J-L**). We observed that new CS+ trials had the highest degree of spatial network synchronization in both early and late reversal (**Fig. S3M**). Interestingly, there was no change in the number of significantly responding astrocytes during either trial type across reversal, while the number of inhibited astrocytes decreased in the ITIs that followed the new CS+-sucrose pairing in expert learners (**Fig. 3R; S3N**). Together, these data suggest there was not a change in astrocytic recruitment across reversal learning, but rather response strengthening and spatial refinement of astrocytic Ca^2+^ dynamics during the new CS+-sucrose pairing as learning stabilized.

To investigate whether astrocytic encoding of task-relevant stimuli was updated during reversal learning, we used our machine learning-based SVM decoder to determine if GCaMP6f fluorescence predicted the old or new sucrose-paired cue, reward delivery, or the old and new CS−specific behavioral responses. In early reversal, astrocytic responses reliably decoded old CS+ trials, incorrect behavioral responses (incorrect licking to the old CS+, incorrect withholding to the new CS+), correct licking to the new CS+, and reward delivery relative to chance. By late reversal learning, astrocytic decoding accuracy significantly improved for new CS+ trials, correct licking to the new CS+, outdated anticipatory licking to the old CS+ (incorrect lick), and reward delivery following the new sucrose-paired cue, while there was no change in the ability of astrocytes to decode old CS+ trials, correct withholding to the old CS+, or incorrect withholding to the new CS+ (**Fig. 3S**). These data strongly indicate that during the initial contingency updating session, astrocytes encode the previously reward-conditioned cue and behavior, as well as appropriate and inappropriate behaviors tied to the new sucrose-paired cue. After animals successfully learned the reward contingency reversal, cortical astrocytes strongly encode the new sucrose-paired cue, updated conditioned behavior, and outdated responses to the old sucrose-paired cue. Collectively, our results establish that astrocytic calcium flexibly tracks updated reward associations, is spatially refined and synchronized across reversal learning, and exhibits learning-dependent encoding of updated, as well as outdated, motivated behavioral action.

### dmPFC astrocytes regulate the magnitude and persistence of conditioned reward seeking

Our results establish dmPFC astrocytes develop new Ca^2+^ responses that track cue-reward associations, rapidly update Ca^2+^ dynamics to reflect new cue-reward contingencies, and display learning-dependent encoding of appropriate and inappropriate behavioral responses. Despite these findings, whether dmPFC astrocytes are required for acquisition of the cue-reward association and expression of conditioned licking was unclear. To address this, we employed the selective gliotoxin, L-AA, to ablate cortical astrocytes during our head-fixed Pavlovian conditioning paradigm. Previous reports demonstrate L-AA-induced astrocytic depletion for up to 6 days following infusion, and that PFC L-AA administration impairs cognitive function, working memory, and reversal learning^22^. Consistent with this, we confirmed that L-AA-induced ablation of dmPFC astrocytes was present 24 hours following injection (**Fig. 4A**). In a separate group of mice, we implanted bilateral guide cannulae aimed at the dmPFC for site-directed infusions of either L-AA or phosphate-buffered saline (PBS) every 6 days throughout Pavlovian conditioning (**Fig. 4B, C**). Mice learned to associate CS+ presentation with sucrose delivery, while no stimuli followed the CS−. All mice acquired the cue-reward association and reliably discriminated between trial types, with no difference in the latency to learn between L-AA- or PBS-treated mice (**Fig. 4D**). Despite both groups learning the CS+-reward association, PBS-treated mice displayed higher rates of anticipatory licking to the CS+ relative to L-AA-treated mice (**Fig. 4E, F**). Neither group showed altered lick rates to the CS− during acquisition (**Fig. 4G**). Accordingly, the number of high-lick trials increased in both groups across conditioning, with PBS-treated mice having a higher percentage of high-lick trials in late learning compared to L-AA-treated mice (**Fig. 4H**). While there were no differences in correct responses between groups in early learning, PBS-treated mice had significantly more correct licking trials and a higher CS+ accuracy in late learning than the L-AA group (**Fig. 4I-K**), suggesting astrocytes modulate the expression of conditioned reward-seeking behavior. There were no differences in CS− correct rate between the groups (**Fig. 4L**), indicating mice appropriately withheld to the non-sucrose paired cue. Taken together, our findings demonstrate that while dmPFC astrocytic ablation does not prevent acquisition of the cue-reward association, it selectively attenuates motivated behavioral action in response to CS+ presentation. These results establish that dmPFC astrocytes functionally regulate the response magnitude of motivated behavior associated with reward-conditioned stimuli.

**Figure 4.**
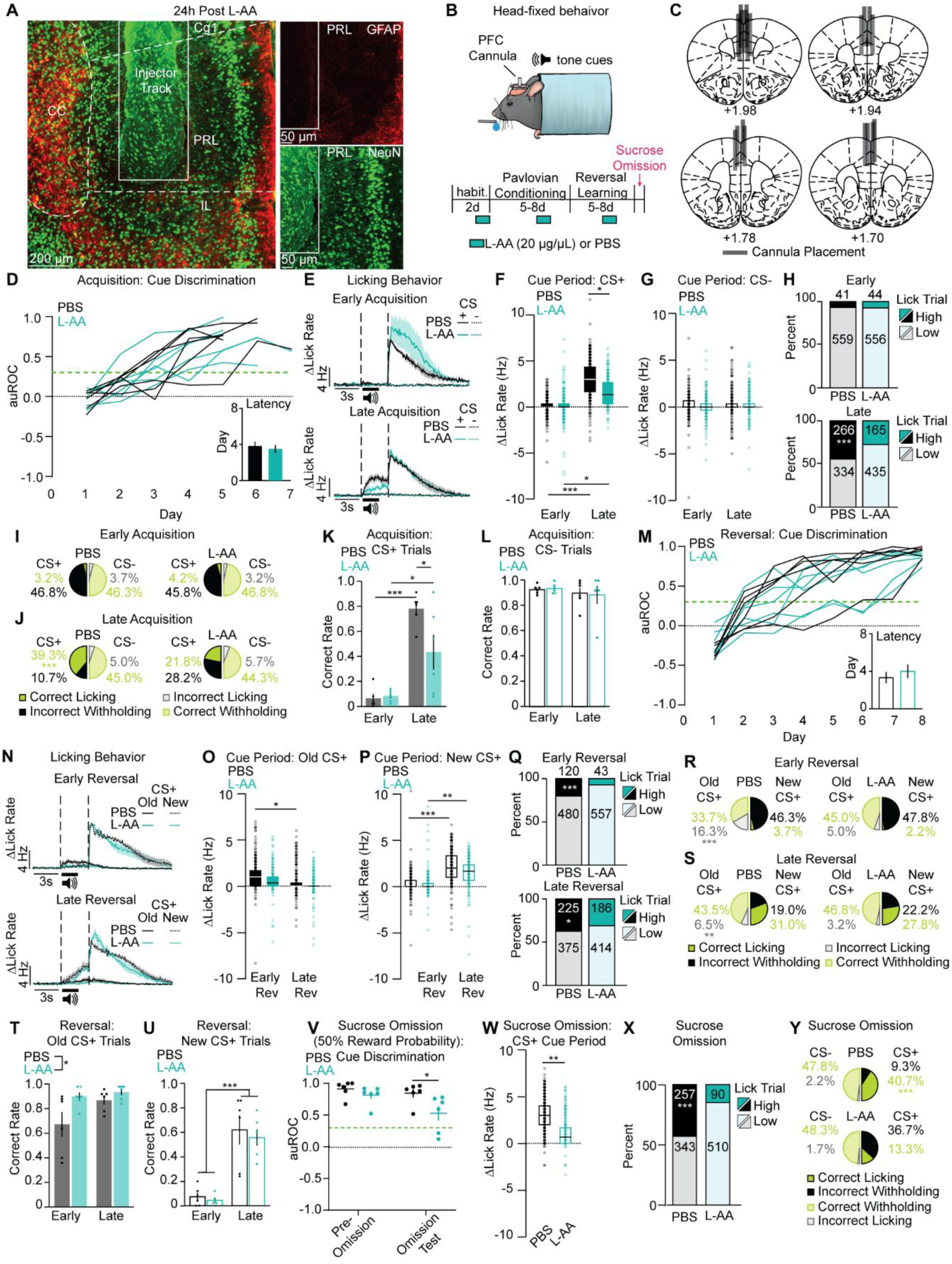
dmPFC astrocytes regulate the magnitude of motivated behavioral action during initial learning, and maintain conditioned seeking when reward contingencies are updated or unpredictable. **A)** Representative 10x (left) and 20x (right) confocal Z-stacks of NeuN (Alexa-488, green) and GFAP (Alexa-594, red) illustrating astrocytic ablation in the dmPFC 24-hours after L-AA injection. **B)** Behavior schematic and experimental timeline. **C)** Representative heatmap of cannula placements along the anterior-posterior axis of the dmPFC. **D)** Behavioral discrimination (anticipatory licking to the CS+ vs CS−) during Pavlovian conditioning across all mice. There was no difference in the latency to acquire conditioned licking to the CS+ between L-AA- and PBS-treated animals (n=6 mice/group; T_10_=0.512, *p*=0.615). Green dashed line represents auROC threshold of 0.3. **E)** Averaged Δlick rate traces show both groups exhibit exclusively consummatory licks in early conditioning (top) and after training both groups display anticipatory licking to the CS+ (late, bottom), with L-AA-treated animals having a less robust conditioned licking response. **F)** There was no difference in the change in anticipatory lick rate during the CS+ cue period between L-AA-treated mice in early learning. Both groups significantly increased the change in anticipatory lick rate to the CS+ across conditioning, with PBS-treated animals having a greater change in anticipatory lick rate during the cue period relative to L-AA-treated mice (Group x Day Interaction: F_1,1196_= 74.75, *p*<0.001). **G)** There were no differences in the change in lick rate to the CS− in L-AA- and PBS-treated mice during early or late learning (Group x Day Interaction: F_1,1184_=0.15, *p*=0.695). **H)** In early learning, both groups had a higher number of low lick trials, with no difference between L-AA- and PBS-treated mice (Χ^2^(1)=0.05, *p*=0.822). While both groups increased the number of high lick trials by late learning (L-AA Early vs Late: Χ^2^(1)=83.43, *p*<0.001; PBS Early vs Late: Χ^2^(1)=219.63, *p*<0.001), L-AA-treated mice had a lower number of high lick trials as compared to PBS-treated mice (Χ^2^(1)=36.21, *p*<0.001). **I)** There were no differences in the behavioral response-type distribution between L-AA- and PBS-treated mice in early learning (CS+: Χ^2^(1)=0.613, *p*=0.434; CS−: Χ^2^(1)=0.105, *p*=0.746). **J)** L-AA-treated mice made significantly less correct responses (high licking in the CS+ cue period) than PBS-treated animals in late learning (Χ^2^(1)=75.89, *p*<0.001), with no differences in responding to the CS− (Χ^2^(1)=0.157, *p*=0.692). **K)** Both groups increased the correct rate of responding to the CS+ across learning, however L-AA-treated mice had a significantly lower CS+ correct rate than PBS-treated mice in late learning (Group x Day Interaction: F_1_,_20_=5.71, *p*<0.05). **L)** There were no differences across learning in the CS− correct rate between L-AA- and PBS-treated mice (Group x Day Interaction: F_1_,_20_=0.07, *p*=0.787). **M)** Behavioral discrimination (anticipatory licking to the new CS+ vs old CS+) during reversal learning across all mice. There was no difference in the latency to acquire conditioned licking to the new CS+ between L-AA- and PBS-treated mice (n=6 mice/group; T_10_=0.698, *p*=0.501). Green dashed line represents auROC threshold of 0.3. **N)** Averaged Δlick rate traces show PBS-treated mice express conditioned licking to the old CS+ upon reward contingency updating in early reversal (top), and both groups display anticipatory licking to the new CS+ by late reversal (bottom). **O)** During reversal learning, L-AA-treated mice did not show differences in the change in lick rate to the old CS+, while PBS-treated animals significantly decreased the change in lick rate from early to late reversal (Group x Day Interaction: F_1,1196_=5.71, *p*<0.05). **P)** Both groups significantly increased the change in anticpatory lick rate during the new CS+ cue period across reversal learning (Group x Day Interaction: F_1,1196_=13.90, *p*<0.001). **Q)** In early reversal, L-AA-treated animals had significantly less high lick trials relative to PBS-treated mice (Χ^2^(1)=41.01, *p*<0.001). While both groups increased the number of high lick trials across reversal learning (L-AA Early vs Late: Χ^2^(1)=108.82, *p*<0.001; PBS Early vs Late: Χ ^2^(1)=44.00, *p*<0.001), L-AA-treated mice had a lower number of high lick trials in late reversal as compared to PBS-treated mice (Χ^2^(1)=5.34, *p*<0.05). **R)** L-AA-treated animals made less incorrect licking errors to the old CS+ than PBS-treated mice upon reward contingency updating in early reversal (Χ^2^(1)=44.58, *p*<0.001), and there were no differences between groups in responding to the new CS+ (Χ^2^(1)=1.942, *p*=0.164). **S)** L-AA-treated animals had less incorrect licking responses to the old CS+ than PBS-treated mice in late reversal (Χ^2^(1)=6.89, *p*<0.01) but did not differ from PBS-treated mice in their behavioral responses to the new CS+ (Χ^2^(1)=2.23, *p*=0.135). **T)** While both groups increased their correct rate to the old CS+, L-AA-treated mice had a higher correct rate than PBS mice during both early and late reversal (Main Effect of Day: F_1,20_=4.15, *p*=0.05; Main Effect of Group: F_1,20_=6.56, *p*<0.05). **U)** Both L-AA- and PBS-treated mice increased in new CS+ correct rate across reversal learning (Main Effect of Day: F_1,20_=56.35, *p*<0.001). **V)** Both groups displayed high cue discrimination prior to the sucrose omission test; however, when the probability of reward was 50% L-AA-treated mice had significantly worse cue discrimination relative to their PBS-treated counterparts (n=6 mice/group; F_1,10_=4.89, *p*=0.05). **W)** During the sucrose omission test, L-AA-treated animals had significantly less of a change in anticipatory lick rate during the CS+ cue period compared to PBS controls (T_10_=3.46, *p*<0.01). **X)** PBS-treated mice had a higher number of high lick trials during the sucrose omission test compared to L-AA-treated animals (Χ^2^(1)=111.72, *p*<0.001). **Y)** L-AA-treated animals had significantly less correct responses (high licking trials during the CS+ cue period) than PBS controls (Χ^2^(1)=178.27, *p*<0.001), with no differences between L-AA- or PBS-treated mice in CS− response types (Χ^2^(1)=0.181, *p*=0.671). L-AA, l-α-aminoadipate; PBS, phosphate buffered saline; auROC, area under the receiver operating characteristic. Bar graphs are represented as mean ± SEM. Post-hoc comparisons: \**p*<0.05, \*\**p*<0.01, \*\*\**p*<0.001.

Next, we examined how L-AA administration affected behavioral adaptation during reversal of the cue-sucrose contingency. Well-trained mice learned that the old CS− (new CS+) now preceded sucrose delivery, while no stimuli followed old CS+ presentation. All mice reversed cue-sucrose association and discriminated between the old and new CS+, with no difference in latency to learn between L-AA- and PBS-treated mice (**Fig. 4M**). During early reversal, PBS-treated mice displayed persistent anticipatory licking to the old CS+ that decreased with reversal learning. In contrast, L-AA-treated mice displayed low lick rates to the old CS+ in early and late reversal (**Fig. 4N, O**). These data suggest that dmPFC astrocytes maintain the expression of previously conditioned behavior during initial reversal of reward contingencies. Both groups displayed comparable changes in lick rate to the new CS+ (**Fig. 4P**), indicating both groups successfully reversed the cue-reward association. PBS-treated mice maintained a greater percentage of high-lick trials compared to L-AA-treated mice across reversal (**Fig. 4Q**). This was reflected in the higher percentage of incorrect withholding trials (anticipatory licking to the old CS+) and lower correct rate to the old CS+ for PBS-treated mice relative to L-AA animals in both early and late reversal (**Fig. 4R-T**). However, both groups increased their new CS+ accuracy with reversal learning (**Fig. 4U**). These results reveal that while both groups successfully updated reward contingencies, PBS-treated mice exhibited persistent conditioned licking to the old CS+, while L-AA mice more readily suppressed responding to the old sucrose-paired cue across reversal learning. Collectively, our data establish that dmPFC astrocytes maintain persistent seeking behavior to previously reward-paired cues despite contingency updating.

Our findings demonstrate L-AA attenuated the magnitude of conditioned licking during initial learning and prevented the perseverance of conditioned reward seeking to the old CS+ upon reversal of reward contingencies. Thus, we examined how L-AA impacted conditioned seeking behavior when there was a 50% probability of sucrose omission following CS+ presentation. During the sucrose omission test, cue discrimination was significantly impaired in L-AA-, but not PBS-treated mice relative to the pre-test session (**Fig. 4V**). Furthermore, during the sucrose omission test, PBS-treated mice had greater CS+-induced changes in lick rate relative to L-AA-treated mice (**Fig. 4W**), indicating a lower rate of anticipatory licks in the L-AA group. Consistently, PBS-treated mice had a greater number of high-lick trials as compared to L-AA mice (**Fig. 4X**), which was reflected in PBS-treated mice having more CS+ correct licking trials than L-AA treated animals (**Fig. 4Y**). These results further support the idea that dmPFC astrocytes modulate the persistence and magnitude of conditioned responses, as L-AA-treated mice began to rapidly extinguish anticipatory licking during times of unpredictable reward. Collectively, our findings demonstrate dmPFC astrocytes regulate the magnitude of motivated behavioral action during initial associative learning and maintain conditioned reward seeking during reversal or unpredictability of reward contingencies.

## DISCUSSION

Here we characterize dmPFC astrocytic Ca^2+^ dynamics during the acquisition, expression, and reversal of cue-reward associative learning, and establish that dmPFC astrocytes regulate the magnitude and persistence of conditioned reward-seeking behavior. We identify that dmPFC astrocytes develop new, learning-dependent Ca^2+^ responses that preferentially encode the cue-sucrose association, as well as correct behavioral action and mistakes during initial associative learning. These Ca^2+^ dynamics rapidly adapt to track updated reward contingencies, and during reversal learning, are strengthened in magnitude and spatially refined to encode the new cue-sucrose association, as well as updated and outdated motivated behavioral action. Finally, we demonstrate that selective ablation of dmPFC astrocytes attenuates motivated behavior during initial cue-reward associative learning and prevents the perseverance of conditioned reward seeking when reward contingencies are updated or unpredictable. Overall, we find that dmPFC astrocytes serve as computational elements that functionally encode cue-reward associations to drive conditioned reward-seeking behavior during initial learning and when contingencies are updated.

These experiments are the first to use 2P microscopy to visualize and longitudinally track the spatiotemporal dynamics of dmPFC astrocytes across various phases of cue-reward associative learning. We found that cortical astrocytes are recruited during Pavlovian conditioning, and their activity patterns are time-locked to behavior, spatially organized, and strongly encode the CS+-sucrose pairing, as well as correct and incorrect behavior. Importantly, spatially organized astrocytic Ca^2+^ dynamics occurred not only during cue presentation but also during ITIs, reflecting engagement beyond sensory or motor epochs. Despite the relatively small number of responding cells, these ITI-associated dynamics suggest a role in feedback integration or outcome evaluation, consistent with astrocytic involvement in ongoing performance monitoring rather than passive stimulus tracking^23^. When we ablated dmPFC astrocytes with the gliotoxin L-AA, we observed decreased anticipatory licking and fewer correct responses to the CS+ as compared to controls, indicating attenuated reward-seeking behavior. Our results are in line with recent work that used fiber photometry and AstroLight tagging to show striatal astrocytic Ca^2+^ tracks learning in an operant cue-sucrose conditioning task, and inhibition of Ca^2+^ in these cells decreased preference for the reward-paired site^30^. Our results and the work of others suggest that astrocytes are regulatory circuit elements that tune motivated behavior and govern the magnitude of reward seeking in response to conditioned cues. Cortical astrocytes differentially respond to excitatory and inhibitory neurotransmission via distinct Ca^2+^ signatures at both the single-cell and network level^34,40^. Glutamatergic activity causes elevated astrocytic Ca^2+^ and increased propagative activity throughout the astrocytic syncytium^34^, which in turn could regulate neuronal synchronization^20^ and long-term potentiation^43,44^. In this context, the enhanced signal strength and temporal synchrony we observed among astrocytes during learning could reflect coordinated prediction updating within the astrocytic network, an emergent computation that may stabilize neuronal ensemble representations of task-relevant cues and outcomes. Previous work has established that dmPFC glutamatergic neurons develop specialized coding patterns that signal the reward-conditioned cue and predict conditioned reward-seeking behavior^12^, and dmPFC output circuits to the ventral striatum and midline thalamus guide conditioned reward seeking through opposing functional activity within these projection-specific populations^5^. Specifically, cortico-striatal projection neurons develop new, excitatory dynamics across Pavlovian sucrose conditioning that are necessary for the expression of conditioned reward seeking, mirroring the astrocytic findings in the current study. Taken with our L-AA results, the learning-dependent elevations in, and spatial organization of, astrocytic Ca^2+^ that we observed positions astrocytes to influence within-dmPFC ensemble formation, functional synaptic plasticity, and circuit-level dynamics that guide reward-motivated behavior.

Overall, our data reveal that dmPFC astrocytic Ca^2+^ dynamics show functional plasticity as behavior updates across distinct learning phases. Across acquisition and reversal of the cue-reward association, dmPFC astrocytic ablation both attenuated the magnitude of anticipatory licking to the CS+ during initial learning, and prevented the persistence of conditioned licking when reward contingencies were updated or reward was unpredictable. Consistently, we found that astrocytic activity strongly encoded the cue-reward pairing and conditioned licking during acquisition, as well as the updated and outdated conditioned behavior during reversal learning. Taken together, our results imply dmPFC astrocytes are key in tuning the response magnitude of conditioned reward seeking when reward-paired cues gain or lose incentive salience^45^ and serve as error detection units when a learned association has become outdated and behavior is inappropriate. These data are consistent with previous findings that L-AA-induced astrocytic ablation in the mPFC impairs attentional set-shifting and reversal learning performance via neuronal loss and dendritic atrophy^22^, suggesting that astrocytes are critical for maintaining the functional integrity of the PFC circuits guiding behavior. Accordingly, astrocytes are preferentially recruited to areas with experience-dependent neuronal plasticity and act to stabilize memory^23^. Given that PFC neuronal activity encodes behavioral response-outcome relationships both within trials and following trial completion, and that disrupting this feedback monitoring impairs behavioral performance^6^, it is likely that dmPFC astrocytic ablation altered cortical neurotransmission, ultimately disrupting feedback monitoring and expression of reward-conditioned behavior. This idea is supported by our evidence that (1) time-locked elevations in dmPFC astrocytic Ca^2+^ encodes both behavioral errors and correct responses across initial learning; (2) learning-dependent astrocytic recruitment in acquisition and cellular response adaptation during reversal; (3) spatial coordination of astrocytic Ca^2+^ dynamics both within trials and subsequent ITIs; and (4) astrocytic ablation attenuated anticipatory licking in initial learning and prevented persistence of conditioned licking when the reward contingency was updated and when there was a 50% chance of reward delivery. We propose that astrocytes utilize these temporally structured Ca^2+^ signals to compute internal feedback predictions and adjust subsequent network states, functionally analogous to prediction error mechanisms described in dopaminergic systems yet implemented through astrocytic signaling. Further, our data reveal that astrocytes strongly encode mistakes across both the acquisition and reversal of Pavlovian sucrose conditioning, and L-AA-treated animals displayed reduced licking and did not attempt alternative behavioral strategies when action-outcome relationships were uncertain. Taken together, our data strongly imply dmPFC astrocytes are key error detection units that critically facilitate neuronal feedback monitoring during reward learning to shape motivated behavior.

The current study employed the selective gliotoxin L-AA to ablate cortical astrocytes *in vivo* and determine the outcome on associative reward learning and reward-conditioned behavior. L-AA is a glutamate analogue that is taken up through sodium-dependent glutamate transporters expressed on astroglia and disrupts the metabolic function of these cells^46–48^, ultimately causing astrocytic ablation and subsequent neuronal damage^22^. Consistent with previous reports that L-AA disrupts PFC-dependent learning and behavioral updating^22^, we show intra-dmPFC L-AA blunted the magnitude of conditioned licking during acquisition, and disrupted the expression of conditioned behavior upon reward contingency updating or during times of unpredictable reward. Given that mice were capable of anticipatory and consummatory licking during both the acquisition and reversal of Pavlovian conditioning, these effects are likely not due to motility issues or the execution of licking behavior. Considering our imaging results establish that astrocytes encode the cue-reward pairing and correct and incorrect behavior across learning, it is likely L-AA disrupts PFC-dependent feedback monitoring, memory updating processes, and weakens the incentive salience of the CS+ that promotes vigorous reward-seeking behavior^6,23,45^. As we repeatedly administered L-AA throughout the current experiment, we could not investigate how astrocytic ablation alters dmPFC neuronal dynamics and encoding across the various stages of Pavlovian conditioning. However, contemporary techniques now allow for more fine-tuned manipulations of astrocytic Ca^2+^ events^21,49–51^. These tools, such as CalEx^50^, can be coupled with *in vivo* 2P imaging strategies to investigate whether decreasing astrocytic Ca^2+^ dynamics alters dmPFC neuronal ensemble formation and maintenance during the acquisition, expression, and reversal of cue-reward conditioning. By combining these more precise astrocytic manipulations with concurrent neuronal Ca^2+^ imaging, the influence of astrocytic activity on the spatiotemporal dynamics of neuronal encoding of task-relevant stimuli, and how this relates to behavioral performance, could be defined. Further, whether driving astrocytic Ca^2+^ during acquisition of cue-reward association improves behavioral performance remains unknown. This is a strong possibility, given that increasing astrocytic Ca^2+^ induces synaptic plasticity, memory improvement, and reward-seeking behavior in the hippocampus and striatum^13,30,31^. The current study provides the foundation for future research to determine precisely how cortical astrocytes integrate into neuronal circuits and whether astrocytic activity influences the neuronal dynamics that guide motivated behavior across reward learning.

Collectively, our results demonstrate that dmPFC astrocytes serve as computational elements that functionally encode cue-reward associations to drive conditioned reward-seeking behavior during initial cue-reward associative learning and when contingencies are updated. We show that cortical astrocytes integrate sensory input, behavioral outcome, and feedback timing to stabilize learning and behavior when response contingencies are initially learned or updated. These data establish a conceptual framework for future studies to investigate how dmPFC astrocytic Ca^2+^ dynamics influence neuronal responding across associative learning, and to examine PFC network dysregulation in both neurons and astrocytes during the progression of neuropsychiatric disorders that disrupt cognitive flexibility^1,32,52–54^. Overall, our findings demonstrate that cortical astrocytes are functionally plastic across reward learning and regulate motivated behavioral action in response to reward-predictive cues.

## Acknowledgements

This study was funded by grants from the National Institute of Drug Abuse (NIDA): F32-DA057794 (JEP), F32-DA053830 (EMD), R01-DA062041, R01-DA051650, & R01-DA054271 (JMO), R01-DA054154 (MDS), R21-DA058901 (JMO & MDS), T32-DA007288 (AMW, RIG, JEP), K99-DA058049 (EMD), the Department of Veteran’s Affairs: VA; I01-BX006179 (JMO), and the MUSC College of Medicine (COMETS; JMO and MDS).

## Author Contributions

JEP, JMO, and MDS designed the experiments and wrote the manuscript. All authors provided technical assistance and intellectual feedback on the project.

## MATERIALS AND METHODS

### Animals

All experiments were approved by the Institutional Animal Care and Use Committee (IACUC) at the Medical University of South Carolina and were conducted in accordance with the NIH-adopted Guide for the Care and Use of Laboratory Animals. Adult male and female C57BL/6J mice (Jackson Laboratory; 8 weeks of age; 20-30 g at start of study) were group housed pre-operatively and single-housed post-operatively on a 12h:12h reverse light:dark cycle (lights off at 8:00am). Mice had access *ad libitum* to standard lab chow, but were water restricted to 90% of initial body weight throughout experimentation. Male and female mice were randomly assigned to experimental groups and experiments were conducted in the dark phase of the light:dark cycle.

### Surgery

For all intracranial surgeries, mice were anesthetized with isoflurane (1-1.5% in oxygen; 1L/minute) and were head-fixed in a stereotaxic frame (Kopf Instruments). Mice were prepared with topical anesthetic (2% Lidocaine; Akorn), 10% povidone, and 70% ethanol at the incision site. Ophthalmic ointment (Akorn), analgesic (Ketorolac 2mg/kg, ip), and 0.9% sterile saline were given pre- and intra-operatively for health and pain management. Antibiotic (Cefazalin 200 mg/kg, sc) was given post-operatively to prevent infection. Mice were given at least one week of post-operative care and recovery.

#### Two-photon calcium imaging surgeries

To target dorsal medial prefrontal cortex (dmPFC) astrocytes for calcium imaging, microinjections of an astrocyte-directed genetically encoded calcium indicator (GECI; AAV5-GfaABC1D-Lck-GCaMP6f, 800 nL; Addgene #52924-AAV5) or an astrocytic GECI mixed 2:1 with a static cytosolic label (AAV5-GfaABC1D-cyto-tdTomato, 800 nL; Addgene #44332) was infused into the dmPFC (AP: +1.85mm; ML: +/− 0.4mm; DV: −2.3 and −2.6mm; 400 nL/DV). A microendoscopic gradient refractive index lens (GRIN lens; 4mm long, 1mm diameter; Inscopix) was then implanted dorsal to the injection site for chronic visual access of prefrontal astrocytes (AP: +1.85mm; ML: +/− 0.4mm; DV: −2.25mm). For head-fixation during experimentation, a custom-made stainless-steel head ring (5 mm ID, 11 mm OD) was secured around the GRIN lens using skull screws and dental cement^12,55,56^. dmPFC GRIN lens placement and GCaMP6f or GCaMP6f/tdT fluorescence was confirmed post-mortem.

#### L-AA validation surgeries

Microinjections of the selective astrocyte toxin L-α-aminoadipate (L-AA; Sigma, cat#A7275) were infused into the dmPFC (AP: +1.85mm; ML: +/− 0.4mm; DV: −2.3 and −2.6mm; 400 nL/hemisphere) and the incision closed using silk sutures. Mice were sacrificed 24-hr post-operative procedure to validate dmPFC astrocytic ablation.

#### Cannulation surgeries

Bilateral double-barrel guide cannula were implanted dorsal to the dmPFC (26-gauge, 8mm length, 3mm below pedestal, 1mm barrel separation; Protech) for site-directed infusions of L-AA. For head-fixed Pavlovian conditioning, a stainless-steel head ring was secured around the cannulae using skull screws and dental cement. Cannula placement was confirmed post-mortem.

### Drug preparation and administration

L-AA was made up fresh daily at a dose of 20 µg/µL^22^. L-AA powder was dissolved in 1N hydrochloric acid, neutralized with 1N sodium hydroxide, and brought to final volume with 0.01 M phosphate buffered saline (PBS; pH = 7.4). As L-AA was previously shown to ablate astrocytes starting at 4-h to 6-d post infusion^22^, mice were given intra-dmPFC microinfusions of L-AA or PBS control through the guide cannula on the last day of habituation (24-h pre-conditioning) and every 6-d during experimentation. To confirm astrocyte ablation, mice received intra-dmPFC L-AA infusions 48-h prior to transcardial perfusion and immunohistochemistry was carried out for cellular markers of astrocytes and neurons.

### Head-fixed Pavlovian sucrose conditioning

#### Acquisition of the tone-sucrose association

Mice were water restricted to 90% of their initial body weight to facilitate cue-reward associative learning and motivated behavior^5,12,57^. Following post-operative recovery, water bottles were removed from the home cage and a watering dish was placed within the cage. Mice were watered once daily with ∼1 mL water, with more or less water given early in experimentation to obtain 90% initial body weight. Mice were monitored and weighed daily to ensure normal health and no dehydration-related issues occurred (as previously^5,12,57^). When weight criteria was reached, mice were habituated for 3-days to head-fixation during 30-minute sessions in which no lickspout, sucrose, or cues were presented. Next, mice underwent cue-sucrose Pavlovian conditioning wherein they were presented with two distinct tone cues (3-kHz pulsing tone, CS−; 12-kHz pure tone, CS+) on a random intertrial interval (ITI; 20-second min, 50-second max) for a total of 100 trials per session (50 CS+/50 CS−). Each tone sounded for 2 seconds and was followed by a 1-second trace interval (TI). Sucrose reward (12.5% sucrose in water; 2.0 µL, <100 µL/session) was delivered via a gravity-driven, solenoid-controlled lick spout following the 12-kHz pure tone and TI, and throughout training became the sucrose-conditioned stimulus (CS+). No reward or stimuli followed the pulsing tone and subsequent TI (CS−). Pavlovian conditioning sessions were separated by at least 24 hours. Cue discrimination was assessed using area under a receiver operator characteristic (auROC) curve to compare the probability that licking during the CS+ and CS− trace intervals was equivalent or significantly higher during CS+ vs. CS− trials after training. auROC values were normalized to be between −1 and 1 by subtracting 0.5 from each auROC value and then multiplying by 2. When mice reached and maintained an auROC value of 0.3 for three or more consecutive days they were classified to be in late learning, as this auROC score confirmed increased anticipatory licking following CS+ presentation (as previously^5,12^). To assess the behavioral responses following each cue type, we classified low and high licking trials below or above threshold of 1.5 Hz ΔLick rate during the cue period (2s tonal cue + 1s TI), and separated the behavioral responses throughout conditioning. Response types were assigned for each trial based on conditioned licking: (1) Correct Licking: high lick trials following the sucrose-predictive cue (CS+); (2) Incorrect Withholding: low lick trials following the sucrose-predictive cue (CS+); (3) Incorrect Licking: high lick trials following non-reward predictive cue (CS−); (4) Correct Withholding: low lick trials following non-reward predictive cue (CS−). For the acquisition imaging experiment, all mice underwent Pavlovian conditioning as described, however, 3 mice did not have two-photon imaging session to record astrocytic activity in early learning, but were included in the early learning behavioral dataset, and these mice had a two-photon imaging session for late learning to record the neural data and time-lock it to behavior.

#### Reversal Learning

After mice were well-trained to execute conditioned licking following the CS+-sucrose pairing, they underwent Pavlovian conditioning sessions in which sucrose delivery occurred following the 3-kHz pulsing tone (old CS−; new CS+) and subsequent trace period. No sucrose delivery followed the pure 12-kHz tone (old CS+). As in acquisition, mice were classified to be in late learning once they reached a licking auROC of above .3, indicative of increased anticipatory licking in the TI following the new CS+. We classified trials as low or high licking based on the 1.5 Hz ΔLick rate threshold during the cue period, and assigned trials to be one of the following behavioral response types based on the contingency updating: (1) Correct Licking: high lick trial following the new sucrose-predictive cue (new CS+); (2) Incorrect Withholding: low lick trials following the new sucrose-predictive cue (new CS+); (3) Incorrect Licking: high lick trials following non-reward predictive cue (old CS+); (4) Correct Withholding: low lick trials following non-reward predictive cue (old CS+). For the reversal learning imaging experiment, one mouse did not acquire the reversal, but it’s behavioral and neural data are included in early reversal as we were interested in this time point as it was the first session of the updated contingency.

#### Omission Testing

To determine if astrocytic activity tracked the cue-reward pairing, or CS+ or reward independently, imaging mice underwent a Sucrose Omission Test and a Cue Omission Test following initial Pavlovian conditioning. During Sucrose Omission, all parameters were consistent with conditioning, however there was a 50% probability that reward would follow CS+ exposure, leading to both CS+ rewarded and unrewarded trials. As during initial learning, no sucrose delivery followed CS− presentation. For Cue Omission, both cues were unplugged and neither the CS+ nor CS− tonal cue was presented. This enabled neural data from imaging experiments to be time-locked across consistent epochs across initial Pavlovian conditioning, reversal learning, and the omission tests. Therefore, in cue omission trials consisted of un-cued sucrose delivery or no stimuli. In both Sucrose and Cue Omission, trials were separated by a variable ITI of 20-50 sec. L-AA- or PBS-treated animals underwent Sucrose Omission testing following acquisition and reversal of Pavlovian sucrose conditioning.

### Two-photon Calcium Imaging

#### Data collection and processing

We visualized GCaMP6f- and tdTomato-expressing dmPFC astrocytes using a two-photon microscope (Bruker Nano Inc) equipped with a tunable InSight DeepSee laser (Spectra Physics, laser set to 920nm, ∼100fs pulse width), resonant scanning mirrors (∼30Hz framerate), a 20X air objective (Olympus, LCPLN20XIR, 0.45NA, 8.3mm working distance), and GaAsP photodetectors. In cases wherein two fields of view (FOVs) were visible through the GRIN lens, we ensured that each FOV was separated by >75 µm in the Z-axis to avoid signal contamination from chromatic aberration and recorded from each FOV during separate imaging sessions. Two-photon imaging was performed during select behavior sessions: Early (days 1-2 of training) and Late Acquisition (after >2 days of at an auROC of .3), Early (days 1-2) and Late Reversal (after >2 days of at an auROC of .3), and during all Omission tests. On each imaging day, we collected a 5-min baseline recording, the behavior session, and 5-min post-behavior recording. To help visualize individual astrocytic boundaries and assist in drawing regions of interest (ROI), we acquired both the static tdTomato and dynamic GCaMP6f during the 5-min baseline and post-session recordings. During the behavioral two-photon sessions, only GCaMP6f was acquired to measure and track astrocytic activity throughout initial reward learning task and contingency updating. All data were acquired with 4-frame averaging using PrarieView software, converted into hdf5 format, motion corrected using SIMA^58^, and all subsequent analyses were performed using custom Python codes in Jupyter Notebook^5,12^ and Visual Studio. A motion-corrected video and averaged time-series frame were used to draw ROIs around dynamic cells using the polygon selection tool in FIJI^59^. An averaged time-series frame of tdTomato and GCaMP6f co-expression from the pre- and post-behavior recording was used to determine individual, visually distinct cells and aid ROI drawing.

### Immunohistochemistry and histology

Mice were deeply anesthetized with isoflurane before transcardial perfusion with ice-cold 0.01 M PBS (pH = 7.4) followed by 4% paraformaldehyde. Brain tissue was extracted for GRIN lens placement and viral expression or cannula placement. Brain tissue was cut into 50 µm coronal sections using a vibratome ahead of immunohistochemistry. Free-floating sections containing the dmPFC were blocked in 0.1M PBS with 2% Triton X-100 (PBST) with 2% normal goat serum (NGS, Jackson Immuno Research) for 2-hours at room temperature with agitation. Sections were then incubated overnight at 4°C with agitation in GFAP (Dako) and/or NeuN (Abcam) primary antisera diluted in 2% PBST with 2% NGS, washed 3 times for 5-minutes in PBST, then incubated in the appropriate secondary antibody (1:1000; Invitrogen) antisera diluted in PBST with 2% NGS for 4-hours at room temperature with agitation. Secondary antibodies were raised in goat and conjugated to Alexa fluorophores. Sections were then washed 3 times for 5-minutes in PBST, mounted on SuperFrost+ slides, and cover slipped with ProLong™ Gold Antifade mounting media. 10x and 20x Z-stacks were collected at 1024×1024 frame size using a Leica SP8 laser-scanning confocal microscope with laser lines that excite at 488 nm and 552 nm.

## QUANTIFICATION AND STATISTICAL ANALYSIS

### Two-photon Calcium Imaging

GCaMP6f fluorescence was recorded during select behavioral sessions and fluorescent traces of each astrocyte was aligned to CS presentation (CS+ or CS−), including the 3-seconds beforehand, 3-seconds between cue onset and sucrose delivery, and 9-seconds after sucrose delivery. For each cue, the 15-second fluorescent trace was averaged across CS+ or CS− trials and plotted as a peri-stimulus time heatmap across astrocytes. To compare astrocytic responses and classify “activated”, “inhibited”, and “non-responding” cells for each session, we calculated the auROC which compared the fluorescence of a 3-second baseline (pre-cue) to the fluorescence surrounding cue presentation, sucrose delivery, and post sucrose consumption (from CS onset to 9-seconds after cue presentation). Each auROC value was normalized to be a value between −1 and 1 by subtracting 0.5 from each auROC and multiplying by 2. Additionally, we quantified and plotted single-cell fluorescent traces from equivalent 15-second time epochs during the inter-trial intervals (ITIs) that followed each trial type. Subsets of astrocytes were reliably identified across behavioral sessions based on shape and relative position within each FOV, allowing us to visualize and longitudinally track response adaptation from individual astrocytes across initial reward learning and contingency updating. Single-cell tracking was performed from Early to Late Acquisition, from Late Acquisition to Sucrose and Cue Omission tests, and from Late Acquisition to Early and Late Reversal Learning. We plotted the auROC scores across each of these behavioral timepoints to evaluate response adaptation across session. Single-cell and tracked auROC scores were analyzed using mixed linear models and omnibus ANOVAs followed by Tukey HSD post-hoc tests to probe significant effects. We also classified and plotted the fluorescence from each trial within this auROC window by behavioral response type and used nested Two-way ANOVAs to compare fluorescent response types followed by Tukey’s HSD post-hoc tests. We then analyzed the relationship between spatial distance and astrocytic response strength (auROC) separately for each trial type and corresponding ITIs. For distance calculations, we computed distances from the cell with the highest auROC to all other cells within each FOV using a direction-weighted metric: Euclidean distance × (1 + 0.2 × cos(θ)), where θ represents the angle from the reference cell. This approach accounts for potential directional biases in cellular responses. We then created scatter plots with pairwise distance from the most active cell on the X-axis and auROC on the Y-axis and calculated the Pearson correlation coefficient to quantify the spatial relationship within both trials and corresponding ITIs.

To assess temporal response similarity within dmPFC astrocytes during each trial type and corresponding ITI, we calculated pairwise cross correlations between the fluorescent traces of individual cells during CS+ trials, CS− trials, and the surrounding ITI of each trial type. Data were separated by trial type and plotted based on cross correlation response similarity. We then calculated each cell’s average correlation coefficient during each trial type and corresponding ITI period. Correlations for CS+ and CS− were compared across behavioral sessions (Early vs. Late Acquisition; Sucrose vs. Cue Omission; Early Reversal vs. Late Reversal) using mixed-effects ANOVAs followed by Tukey HSD post-hoc comparisons. Additionally, pairwise correlations between cells were used to determine spatial response similarity between nearest neighbors within FOVs during each trial type and ITI. We used each cell as a central reference point, mapped the corresponding correlations onto the other cells within each FOV, and repeated this for each cell within each FOV for CS+ trials, CS− trials, and ITI periods. These spatial plots were accumulated across all cells in our dataset and we mapped the average correlation as a function of distance and position away from the central reference cell. To account for edge cases, we calculated the average cell diameter across our dataset and excluded spatial data that was 11 cell radii (zones) from the reference cell, and we quantified distance-correlation data within each zone. To isolate trial-specific spatial relationships between cells, we calculated and plotted the difference in average correlation coefficient for each cell during each trial type relative to the respective ITI period. Data was analyzed using Mixed Linear Models and Two-way ANOVAs followed by Tukey’s post-hoc comparisons.

We used binary decoding to determine if the activity of each astrocyte predicted task-relevant stimuli such as trial type (CS+ or CS−), reward delivery, or behavioral response type (Correct Licking, Correct Withholding, Incorrect Licking, Incorrect Withholding). We implemented machine-learning-based trial-level temporal decoding using Scikit-learn’s SVM classifier (SVC), *Sklearn.svm.SVC,* that was informed by the fluorescence of each astrocyte during a baseline and event epoch. The baseline period was the preceding 3-second period before cue presentation and the event epoch varied for each decoding stimulus to capture the relevant period: CS+ and CS− as the 3-second period after CS onset (tonal cue + TI), reward as the 3-second period following sucrose delivery (or omission during Sucrose Omission wherein there was 50% probability of sucrose reward), and the behavioral response types as a 3-second period from the trace interval through sucrose delivery. We used Leave-One-Out Cross-Validation (LOOCV) for our decoding analysis. As a control, the labels randomly shuffled 10 times per cell, and the decoding analysis was repeated. To normalize decoding scores, we used a per-cell normalization in which the shuffled data from each astrocyte was subtracted from the real data for that cell. Normalized decoding scores were compared within and between behavioral sessions using Mixed Linear Models and omnibus ANOVAs followed by Tukey’s post-hoc tests.

### Behavioral Data

Licking data was analyzed by calculating the ΔLick rate in Hertz (Hz) during the 3-second cue period (2-second cue + 1-second TI) or the 3-seconds following reward delivery as compared to a 3-second pre-cue baseline. All ΔLick rate data was analyzed across behavioral sessions and groups using nested Mixed Linear Models and omnibus ANOVAs followed by Tukey’s post-hoc tests. For the sucrose omission test of L-AA and PBS-treated mice, the change in lick rate was analyzed by unpaired t-Test of the mean trial ΔLick rate per animal. auROC of licking behavior for the imaging experiments were compared across behavioral sessions using paired t-Tests, and in the L-AA experiment using an unpaired t-Test between the experimental and control group. High and low licking trials were compared across behavioral sessions and groups using Chi-Square tests. Behavioral response types (Correct Licking, Correct Withholding, Incorrect Licking, Incorrect Withholding) were compared across behavioral sessions and groups using Chi-Square tests. CS+ Correct Rate and CS− Correct Rate were analyzed using two-way ANOVA and Tukey’s post-hoc comparisons. All statistical analyses were performed in GraphPad Prism statistical software or Python.

## Supplementary Information

**Figure S1 (Related to Figure 1).**
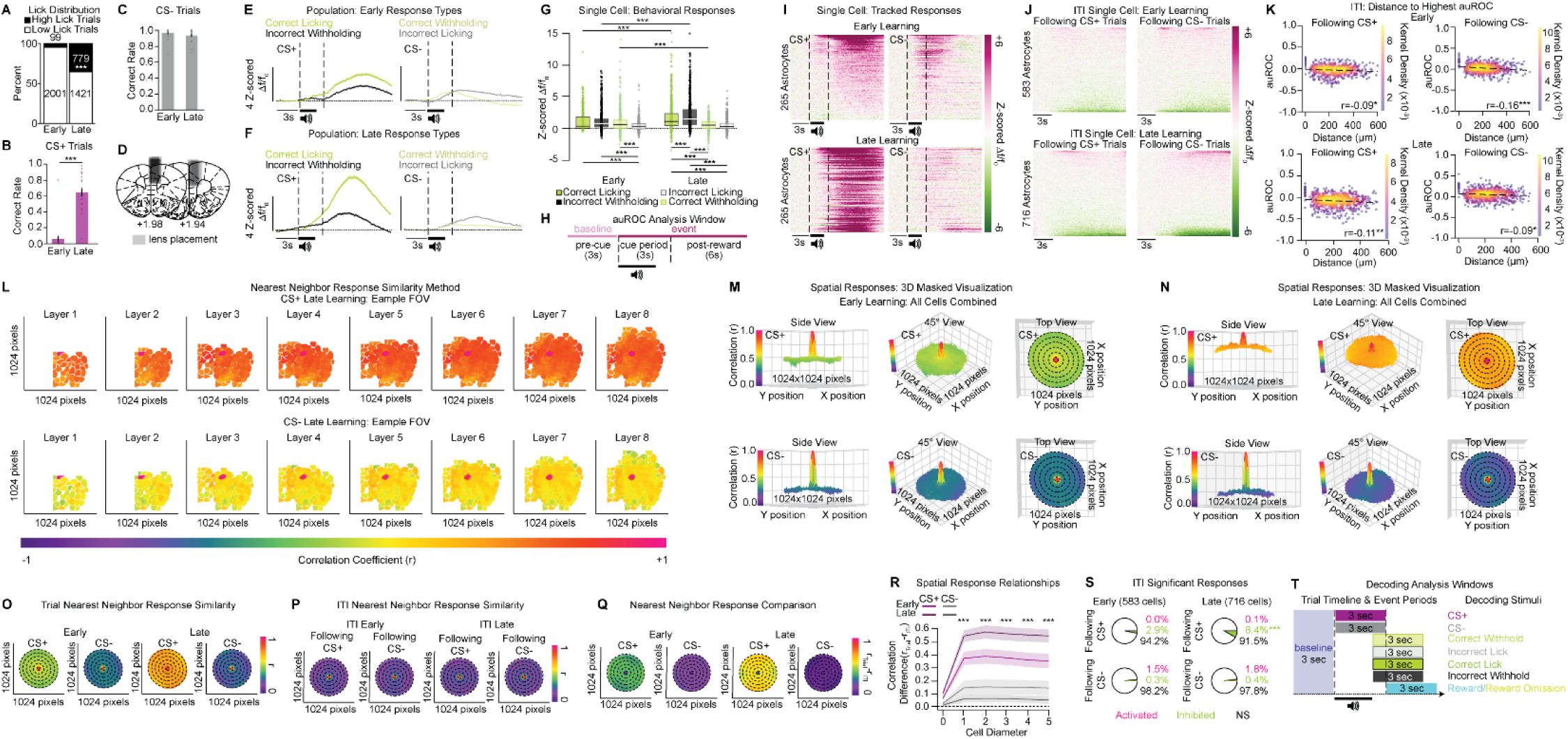
dmPFC astrocytic trial and ITI response patterns across Pavlovian sucrose conditioning. **A)** The number of high lick trials significantly increased from early to late learning (Χ^2^(1)=621.08, *p*<0.001). **B)** The CS+ correct rate significantly increased across Pavlovian sucrose conditioning (n=20 mice, 21-22 FOVs; T_41_=-9.532, *p*<0.001). **C)** There was no difference in CS− correct rate between early and late learning (T_41_=1.406, *p*=0.167). **D)** Representative heatmap of lens placements along the anterior-posterior axis of the dmPFC. **E, F)** Averaged traces from early (**E**) and late (**F**) conditioning show the population activity during CS+ and CS− trials separated by behavioral response type (Early: n=17 mice, 17 FOVs; 583 cells; Late: n=20 mice, 22 FOVs, 716 cells). **G)** In early learning, the fluorescent responses were greatest during correct licking (high licking to the CS+), incorrect withholding (low licking to the CS+), and correct withholding (low licking to the CS−) trials relative to incorrect licking trials (high licking to the CS−). In late conditioning sessions, astrocytic fluorescent signal increased during correct licking and incorrect withholding to the CS+ and decreased during correct withholding to the CS− relative to early learning, with incorrect withholding to the CS+ have the greatest signal in late learning (Response Type x Day Interaction: F_3, 4696_=39.81, *p*<0.001). **H)** auROC analysis window used to analyze astrocytic GCaMP6f fluorescence during behavioral 2P imaging sessions. **I)** Single-cell heatmaps of tracked cells between early and late learning sessions depict response adaptation across learning, with biased activation towards the CS+-sucrose pairing (n=265 cells). **J)** Single-cell heatmaps of astrocytic responses during inter-trial intervals (ITIs) that follow CS+ or CS− trials in early (top; n=583 cells) and late learning (bottom; n=716 cells) show mild patterns of activation and inhibition. **K)** Direction-weighted distance correlations of auROC values from ITIs following CS+ and CS− across learning. auROC values for ITIs that were preceded by CS+ and CS− trails were significantly correlated with distance in early (Pearson-R value on each graph; CS+ ITI *p*<0.05; CS− ITI *p*<0.001) and late conditioning (Pearson-R value on each graph; CS+ *p*<0.01; CS− *p*<0.05). **L)** Spatial response synchrony mapping method for each trial type. Each cell within an FOV was used as a central reference point and remaining cells within that FOV were assigned corresponding correlations of fluorescent signal relative to the reference cell. Each new reference cell created a new layer with neighboring cells mapped spatially relative to the reference cell onto a 1024×1024 grid. **M, N)** 3-dimensional spatial correlation plots were created that combined all cells in CS+ and CS− trials in early (**M**) and late (**N**) learning. To account for edge cases, the average cell diameter across the dataset was calculated and masked to 11 cell radii (5-cell zones), with cells outside of this spatial distance excluded from analysis. Distance-correlation data was calculated within each zone during CS+ and CS− trials, as well as corresponding ITIs. **O-Q)** Spatial response maps from each trial type (CS+, CS−; **O**), subsequent corresponding ITI (**P**), and the difference between the two (**Q**) in early and late learning depict response correlations within a 5-cell distance. Dashed lines correspond to a 1-cell diameter distance zone. **R)** Astrocytes have the highest spatial response similarity during normalized CS+ trials in late learning across all distance zones as compared to early normalized CS+ trials and CS− trials from both early and late conditioning (Day x Trial type interaction: F_1,456_=45.087, *p*<0.001). **S)** There was an increase in significantly inhibited cells during ITIs following CS+ trials across learning (Fisher’s Exact Test, *p*<0.001), with no changes in ITI activity following CS− trials from early to late training (Fisher’s Exact Test, *p*=0.943). **T)** Time epochs used in the decoding analysis for each decoding stimulus. auROC, area under the receiver operating characteristic; ITI, inter-trial interval. Post-hoc comparisons: \**p*<0.05, \*\**p*<0.01, \*\*\**p*<0.001.

**Figure S2 (Related to Figure 2).**
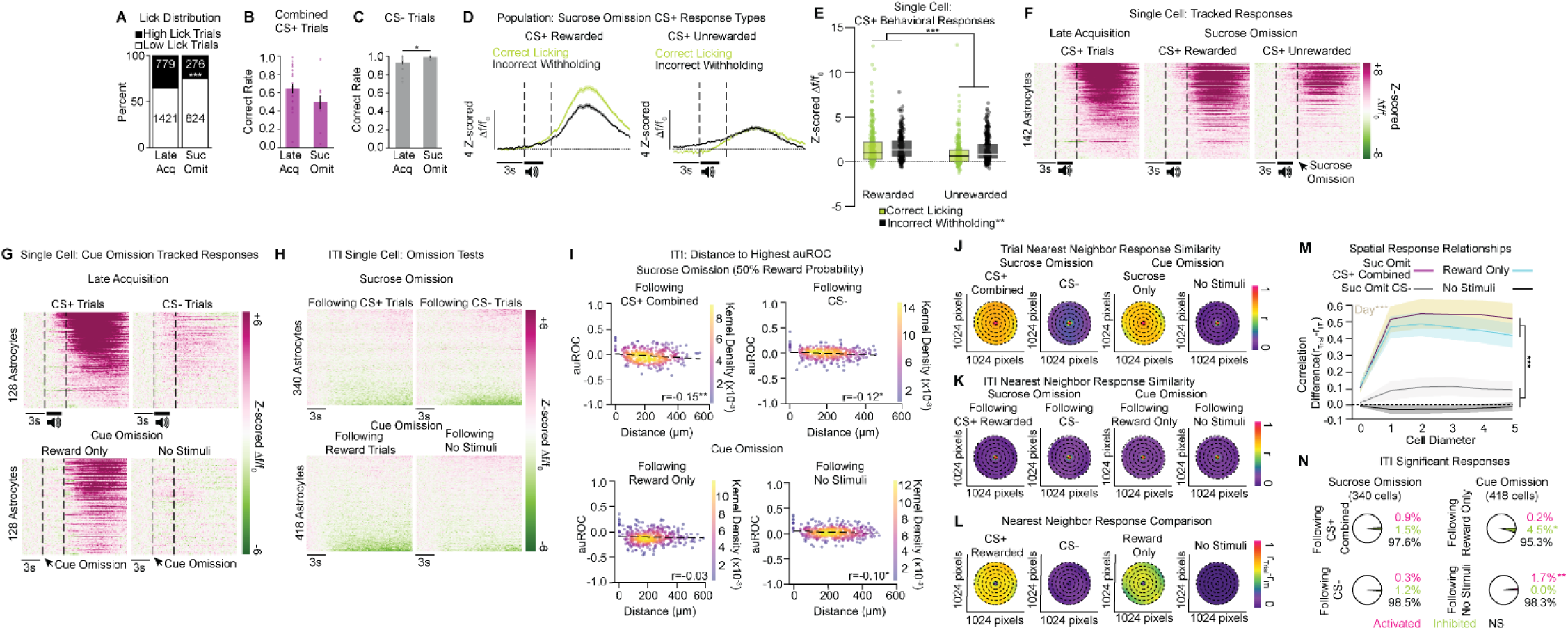
dmPFC astrocytic trial and ITI response patterns during the sucrose omission (50% reward probability) and cue omission tests. **A)** The percent of high licking trials significantly decreased from late learning when there was a 50% probability of reward (Χ^2^(1)=35.42, *p*<0.001). **B, C)** There was no difference in the CS+ correct rate from late acquisition to the sucrose omission test (**B;** T_31_=1.74, *p*=0.092), while CS− correct rate significantly increased (**C;** T_31_=-2.46, *p*<0.05). **D)** Averaged traces from CS+ rewarded and unrewarded trials show the population activity separated by correct and incorrect responses during 50% reward probability in the sucrose omission test (n=11 mice, 11 FOVs, 340 cells). **E)** Astrocytic Ca^2+^ elevations were strongest during rewarded trials than when sucrose was omitted, and cells were most responsive when mice incorrectly withheld licking to the CS+ (Main Effect of Reward: F_1,1286_=29.39, *p*<0.001; Main Effect of Behavioral Response: F_1,1286_=7.42, *p*<0.01). **F)** Single-cell heatmaps of tracked cells between CS+ in late learning to CS+ rewarded and unrewarded trials show astrocytes display the greatest response patterns to the CS+-reward pairing (n=142 cells). **G)** Single-cell heatmaps of tracked cells between late learning to the cue omission test demonstrate astrocytic Ca^2+^ dynamics are strongest when CS+ predicts reward delivery then when sucrose is presented alone (n=128 cells). **H)** Single-cell heatmaps of astrocytic responses during ITIs following CS+ and CS− trials during sucrose omission (top, n=340 cells) and Reward Only or No Stimuli trials during cue omission (bottom, n=418 cells) show mild patterns of activation and inhibition. **I)** Direction-weighted distance correlations of auROC values from ITIs surrounding trials in the sucrose and cue omission tests. During sucrose omission there were response-distance relationships in the ITIs following CS+ trials combined and CS− trials (Pearson-R value on each graph; CS+ combined ITI: *p*<0.01; CS− ITI: *p*<0.05). Conversely, during cue omission there was not a spatial distance-response relationship in the ITIs following Reward Only trials (Pearson-R value on graph; Reward Only ITI: *p*=0.48, yet one was present following trials with No Stimuli (Pearson-R value on graph; No Stimuli ITI: *p*<0.05). **J-K)** Spatial response maps from each trial type (CS+ combined, CS−, Reward Only, No Stimuli) in the omission tests (**J**), subsequent corresponding ITI (**K**), and the difference between the two (**L**) depict the nearest neighbor response correlations within a 5-cell distance. Dashed lines correspond to a 1-cell diameter distance zone. **M)** There were no differences in spatial response similarity between normalized sucrose omission CS+ trials combined and cue omission Reward Only trials, with both having greater spatial response synchronization than normalized CS− or No Stimuli trials (Main Effect of Trial Type: F_1, 252_=287.267, *p*<0.001). Overall, normalized trials in sucrose omission had greater response synchrony than normalized cue omission trials (Main Effect of Day: F_1, 252_=12.384, *p*<0.001). **N)** The number of significantly inhibited astrocytes increased in the ITIs surrounding Reward Only trials relative to CS+ trials when reward probability was 50% (Fisher’s Exact Test, *p*<0.05). There was an increase in activated astrocytes in the ITIs following No Stimuli trials relative to CS− trials in sucrose omission (Fisher’s Exact Test, *p*<0.001). auROC, area under the receiver operating characteristic; ITI, inter-trial interval. Post-hoc comparisons: \**p*<0.05, \*\**p*<0.01, \*\*\**p*<0.001.

**Figure S3 (Related to Figure 3).**
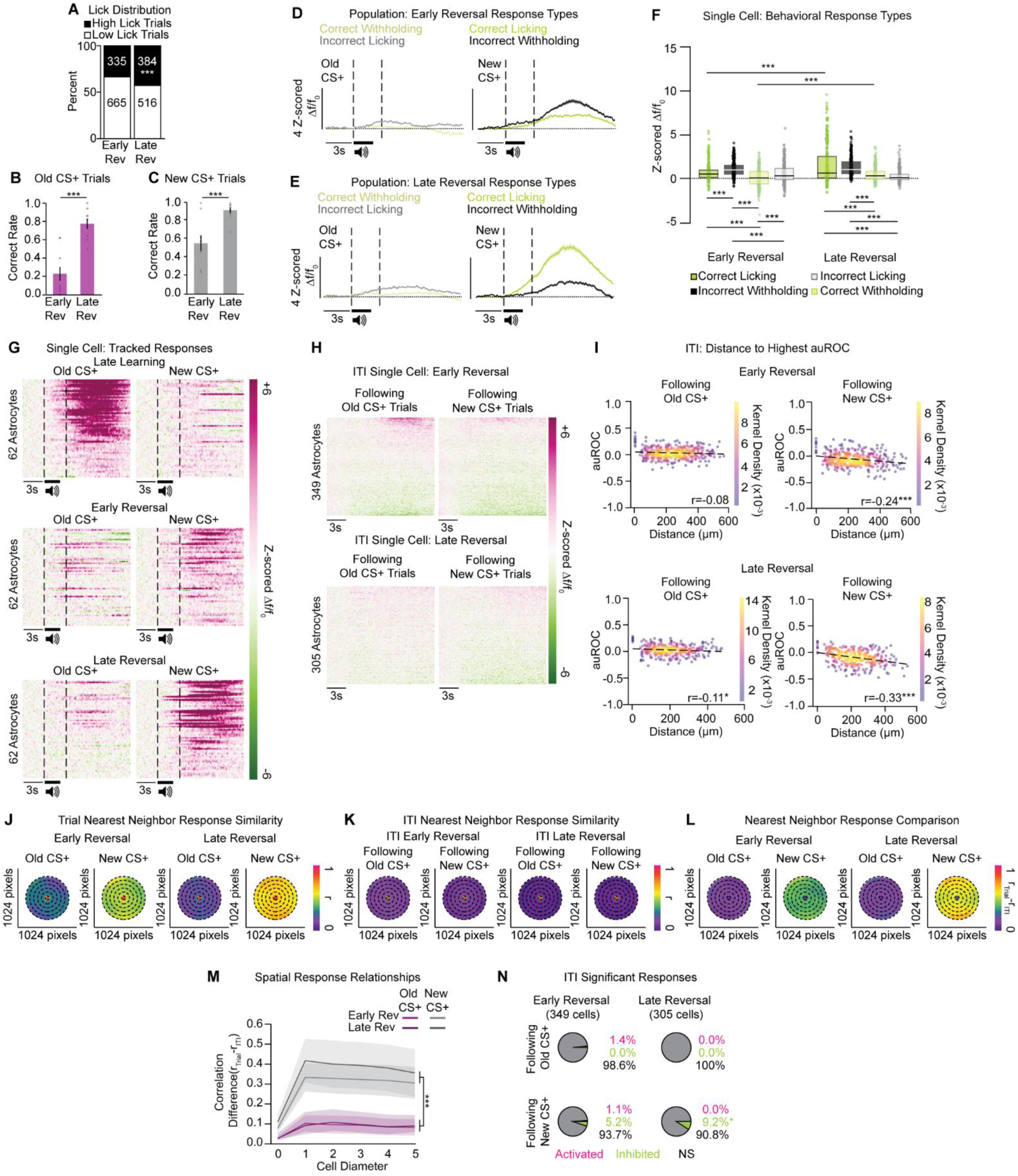
dmPFC astrocytic trial and ITI response patterns during contingency updating and reversal learning. **A)** The number of high lick trials increased across reversal learning (Χ^2^(1)=16.53, *p*<0.001). **B, C)** Mice learned the updated reward contingency and increased correct rate of responding for the old (**B**) and new (**C**) CS+ (n= 8-9 mice; old CS+: T_17_=-5.78, *p*<0.001; new CS+: T_17_=-3.83, *p*<0.001). **D, E)** Averaged traces from early (**D**) and late (**E**) reversal show the population activity during new and old CS+ trials separated by behavioral response type (Early Rev: n=9 mice, 10 FOVs, 349 cells; Late Rev: n=8 mice, 9 FOVs, 305 cells). **F)** Upon updating to the new CS+-sucrose pairing in early reversal, astrocytic fluorescent responses were greatest during behavioral errors (incorrect withholding to the new CS+, followed by incorrectly licking to the old CS+), as fluorescent signal during correct licking to the new CS+ and correct withholding to the old CS+ increased across reversal learning. In late reversal, astrocytic responses were greatest during correct licking and incorrect withholding to the new CS+ relative to behavioral responses to the old CS+ (Response Type x Day Interaction: F_3,2462_=24.76, *p*<0.001). **G)** Single-cell heatmaps of tracked cells between late Pavlovian conditioning and reversal learning show rapid adaptation in astrocytic response patterns shift from the old CS+-sucrose association to the new CS+-sucrose pairing (n=62 cells). **H)** Single-cell heatmaps of astrocytic responses during ITIs following the old and new CS+ trials during early (top, n=349 cells) and late (bottom, n=305 cells) reversal show mild response dynamics. **I)** Direction-weighted distance correlations of auROC values from ITIs surrounding old and new CS+ trials across reversal learning. In early reversal, there was a selective spatial distance-response relationship in the ITIs surrounding the new CS+ (Pearson-R value on each graph; old CS+: *p*=0.120; new CS+: *p*<0.001). In late reversal, ITIs following both the and new CS+ had a significant auROC-distance correlations (Pearson-R on each graph; old CS+ ITI: *p*<0.05; new CS+ ITI: *p*<0.001). **J-L)** Spatial response maps from old and new CS+ trials across reversal learning (**J**), subsequent corresponding ITIs (**K**), and the difference between the two (**L**) show the nearest neighbor response correlations within a 5-cell distance. Dashed lines correspond to a 1-cell diameter distance zone. **M)** Across reversal learning, astrocytes have the highest spatial response similarity during normalized new CS+ trials relative to normalized old CS+ trials (Main effect of Trial type: F_1, 204_=64.958, *p*<0.001). **N)** The number of significantly responding cells did not change from early to late reversal in the ITIs following the old CS+ (Fisher’s Exact Test, *p*=0.064). Across reversal learning, the number of significantly inhibited astrocytes increased in the ITIs following new CS+ trials (Fisher’s Exact Test, *p*<0.05). auROC, area under the receiver operating characteristic; ITI, inter-trial interval. Post-hoc comparisons: \**p*<0.05, \*\**p*<0.01, \*\*\**p*<0.001.

